# Competitive Olivocerebellar Input Selection Promotes Resilient Circuit Formation

**DOI:** 10.64898/2026.06.16.732615

**Authors:** Jonathan A Coello, Kristen M Crane, Alyssa M Lyon, Annie E Walls, Presley A Pickeral, Jack R Fernandez, Breanne L Dao, Ethan A Bongiovanni, Austin L Fitzgerald, Meike E van der Heijden

## Abstract

Many neural circuits undergo competitive input selection, a process in which supernumerary connections compete for innervation territory on target cells. This process can create atypical circuits when functional inputs are favored over compromised ones. Yet, it is often unclear whether such atypical circuits are maladaptive or promote functional resilience. We investigated this using the olivocerebellar climbing fiber circuit, where multiple inputs compete to mono-innervate Purkinje cells. We found that eliminating neurotransmission from ∼50% of olivocerebellar neurons reduced climbing fibers’ competitiveness during input selection and decreased survival of parental inferior olive neurons. Conversely, functional climbing fibers expanded their innervation territory. Despite these atypical circuits, climbing-fiber-dependent motor control was only minimally affected and social behaviors were fully preserved. These results demonstrate that neurotransmission-dependent competition promotes resilient circuits formation, maintaining complex behaviors even when a large proportion of inputs to the cerebellum are developmentally compromised.

## INTRODUCTION

Transient supernumerary connections are a common feature of developing neural circuits and are often refined through competitive selection among inputs of varying strength.^1–4^ In sensory systems, competitive synapse selection is essential for establishing precise topographic maps,^1,5,6^ and experimentally disrupting this process can impair circuit specificity and function. However, developmental redundancy may also promote resilient circuit assembly when multiple wiring patterns can support largely normal behavioral outcomes.^7,8^ It is often difficult to disentangle whether atypical wiring patterns cause maladaptive behavioral control or present a mechanism for resilient circuit assembly, requiring precise experiments to determine the behavioral relevance of structural changes.

Here, we use the highly tractable olivocerebellar circuit to investigate how competitive input selection contributes to adaptive circuit development. Specifically, we examine how compromising a large subset of neural inputs during development influences circuit assembly and the acquisition of complex behaviors.

The olivocerebellar system is particularly well suited for studying the functional relevance of competitive input selection because each cerebellar Purkinje cell is initially innervated by multiple climbing fibers, but only a single climbing fiber is ultimately retained in adulthood.^9,10^ This developmental refinement is thought to depend on neural activity. Neuroanatomical and electrophysiological studies demonstrate a progressive reduction in the number, size, and strength of weaker climbing fiber inputs, accompanied by strengthening of dominant inputs.^11,12^ Repeated climbing fiber activation further reinforces strong synapses while weakening weaker competitors, that are likely to be eliminated.^13,14^ Consequently, reducing neurotransmission in a subset of climbing fibers should diminish their competitive advantage and alter the final pattern of olivocerebellar connectivity.

Olivocerebellar climbing fibers are also critical for cerebellar function and behavior.^15^ Climbing fibers transmit error signals required for motor learning and have also been implicated in cognitive and social behaviors.^16^ Disrupted climbing fiber signaling is associated with several movement disorders: widespread climbing fiber miswiring causes ataxia,^17^ loss of climbing fiber neurotransmission produces severe dystonia,^18^ and excessive climbing fiber activity induces tremor.^19,20^ Abnormal climbing fiber innervation and impaired input elimination have also been reported in mouse models of Down syndrome,^21^ autism spectrum disorders,^22,23^ and schizophrenia.^24^ Together, these findings highlight the importance of climbing fiber development for a broad range of motor and non-motor behaviors.^25,26^

To determine how altered climbing fiber selection affects behavioral outcomes, we used *Pdx1^Cre^-*mediated recombination to selectively label and compromise a large subset of olivocerebellar climbing fibers during development. This manipulation produces atypical climbing fiber wiring because functional inputs are preferentially selected during competitive selection. Despite substantial developmental alterations in olivocerebellar connectivity, we found that most climbing-fiber-dependent motor and social behaviors remained intact in both juvenile and adult mice. These findings demonstrate that competitive input selection promotes resilient olivocerebellar circuit assembly and enables robust behavioral development even when a large proportion of climbing fiber inputs is developmentally compromised.

## RESULTS

### *Pdx1^Cre^* maps a subset of climbing fibers throughout the olivocerebellar circuitry

To investigate how competitive climbing fiber selection influences behavioral outcomes, we leveraged a genetic strategy that provides access to a subset of climbing fibers throughout the developing cerebellum. Previous studies reported expression of the *Pdx1* promoter in some inferior olive neurons that give rise to climbing fibers innervating the cerebellar cortex.^27,28^ Here, we demonstrate that *Pdx1^Cre^*possesses properties ideally suited for mapping and manipulating the developing olivocerebellar circuit to induce atypical circuit formation.

To visualize *Pdx1^Cre^*-expressing neurons and their projections, we crossed *Pdx1^Cre^* mice with a Cre-dependent tdTomato reporter line (Figure 1A). *Pdx1^Cre^*-negative and *Pdx1^Cre^*-positive climbing fibers could be distinguished by the absence or presence of tdTomato labeling, respectively (Figure 1B). Because the presynaptic marker vGluT2 selectively labels climbing fibers within the molecular layer of the cerebellar cortex, where Purkinje cell dendrites reside, it allowed us to visualize the full climbing fiber population.

**Fig 1.**
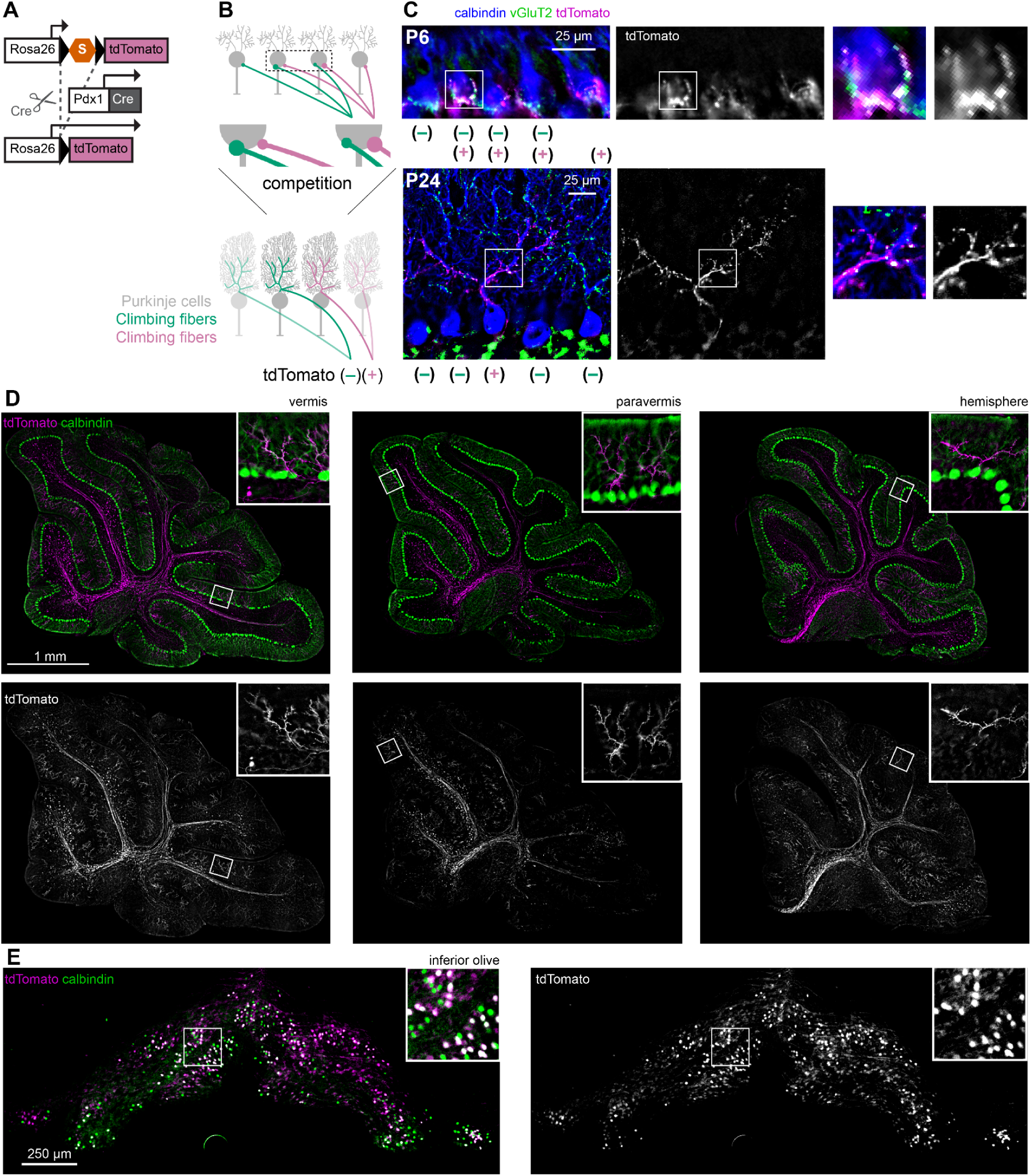
Selectively labeling *Pdx1^Cre/+^* neurons with tdTomato reveals competition amongst tdTomato(+) and tdTomato(–) climbing fibers. **A.** Genetic strategy that expresses the tdTomato reporter in all neurons upon *Pdx1^Cre^-*mediated recombinase. **B.** Schematic illustrates tdTomato(+) and tdTomato(–) climbing fibers initially multi-innervating Purkinje cell somas and compete for mono-innervation of the dendrites. **C.** Top row shows at postnatal day (P) 6 both tdTomato(+) and tdTomato(–) climbing fibers are found in the Purkinje cell bodies. Bottom row shows at P24 Purkinje cells are mono-innervated by tdTomato(+) or tdTomato(–) climbing fibers on dendrites. Top inset: multi-innervation on Purkinje cell body. Bottom inset: mono-innervation on Purkinje cell dendrite. Blue = calbindin (Purkinje cells); Green = vGluT2; Purple = tdTomato (also in white, right). **D.** Sagittal cerebellar sections from the vermis to the hemisphere (medial to lateral). Insets show tdTomato(+) climbing fibers present throughout the cerebellar cortex. Green = calbindin (Purkinje cells); Purple = tdTomato (also in white, bottom row). **E.** Mosaic pattern of tdTomato(+) and tdTomato(–) inferior olive neurons. Inset shows some, but not all, inferior olive neurons are tdTomato(+). Green = calbindin (inferior olive neurons); Purple = tdTomato (also in white, right). Images are representative for N = 3 mice.

At postnatal day 6 (P6), before climbing fiber selection is complete, tdTomato(+) and tdTomato(–) climbing fibers were frequently observed to converge onto the same Purkinje cells (Figure 1C, top row). By P24, when climbing fiber selection is complete, some Purkinje cells were mono-innervated by tdTomato-positive climbing fibers (Figure 1C, bottom row), demonstrating that unmanipulated *Pdx1^Cre^*-positive climbing fibers can successfully compete for and be selected as the sole climbing fiber input. Together, these findings indicate that *Pdx1^Cre^* labels a subset of climbing fibers that participates in normal competitive input selection alongside unlabeled climbing fibers.

We next assessed the spatial distribution of tdTomato(+) climbing fibers throughout the cerebellum. tdTomato(+) fibers were present across the rostrocaudal extent of the cerebellar cortex and throughout the vermis, paravermis, and hemispheres (Figure 1D). Similar widespread labeling was observed in coronal sections (Supplemental Figure 1). These results demonstrate that *Pdx1^Cre^* labels a broadly distributed subset of climbing fibers that innervates Purkinje cells throughout the cerebellar cortex.

Finally, we examined the distribution of labeled neurons within the inferior olive. Consistent with our observations in the cerebellar cortex, tdTomato(+) and tdTomato(–) neurons were intermingled throughout the inferior olive without obvious regional segregation (Figure 1E). Thus, *Pdx1^Cre^* labels a subset of inferior olive neurons independent of anatomical subregion.

Collectively, these findings establish Pdx1Cre as a powerful genetic tool for targeting a broadly distributed subset of olivocerebellar climbing fibers and studying how competitive input selection shapes circuit development and behavioral function.

### Neurotransmission promotes climbing fibers’ competitiveness during input selection

Next, we investigated whether disrupting neurotransmission in *Pdx1^Cre^*-positive climbing fibers would reduce their competitiveness during climbing fiber selection. To do this, we conditionally deleted the gene encoding vGluT2 (Figure 2A), the vesicular glutamate transporter responsible for packaging glutamate into presynaptic vesicles. In the absence of vGluT2, synaptic vesicles cannot be loaded with glutamate, preventing *Pdx1^Cre^*-positive neurons from transmitting signals to postsynaptic Purkinje cells (Figure 2B).

**Figure 2.**
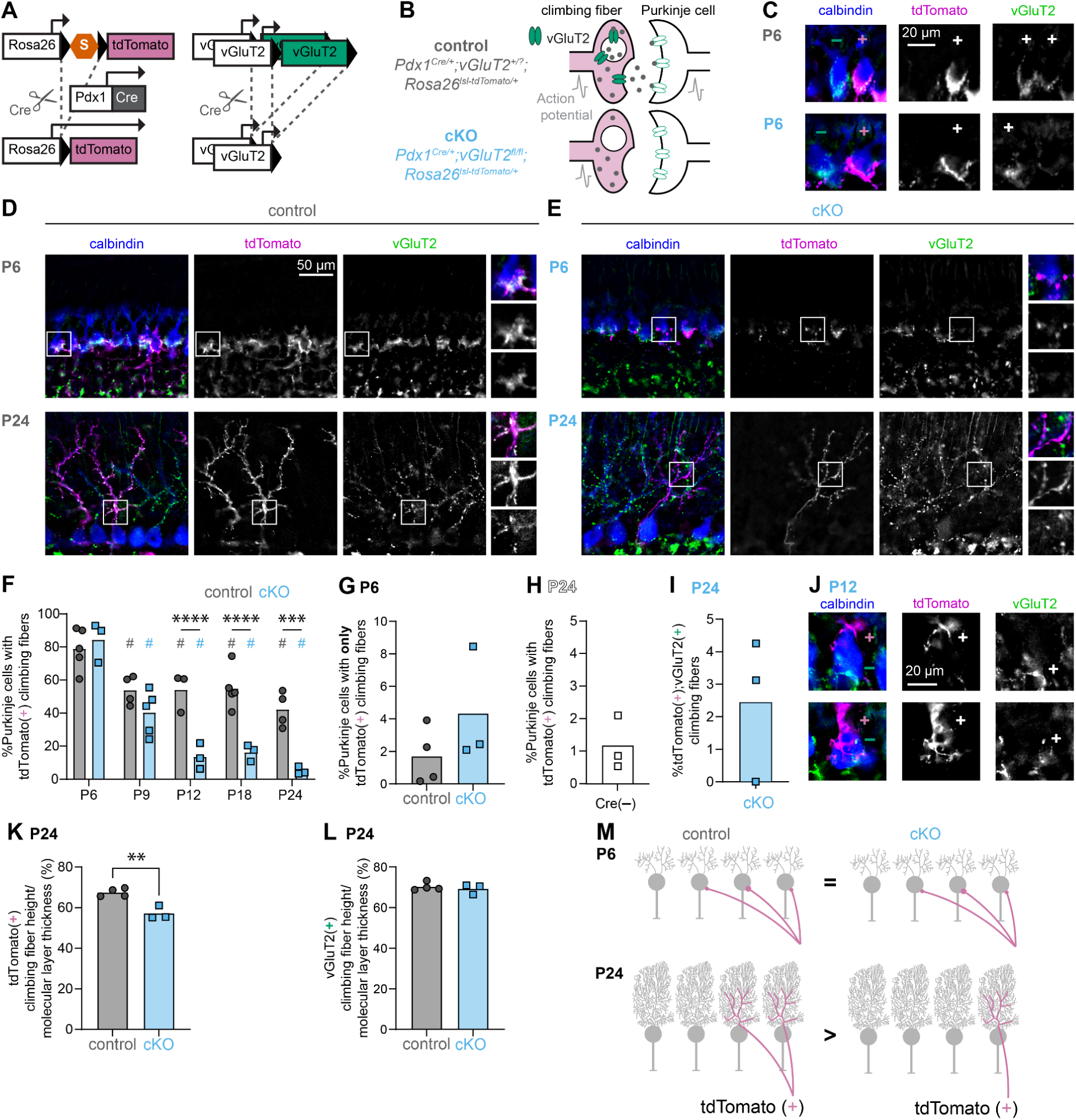
Conditional deletion of vGluT2 from the Pdx1Cre/+ climbing fibers leads to excessive tdTomato(+) climbing fiber elimination after P12. **A.** Genetic strategy used to generate a conditional knock-out model in which the tdTomato reporter is expressed in all *Pdx1^Cre^*-positive neurons concomitant with selective vGluT2 from these neurons. **B.** Schematic showing how conditional deletion of vGluT2 only affects fast neurotransmission in Pdx1Cre/+ climbing fibers when both copies of vGluT2 are floxed out. **C.** At P6, tdTomato(+) climbing fibers are observed in the Purkinje cell somas in both control and cKO cerebella. In cKO cerebella, these tdTomato(+) climbing fibers lack detectable vGluT2 signal. Blue = calbindin (Purkinje cells); Purple = tdTomato; Green = vGluT2. **D.** Representative images from control mice. At P6, tdTomato(+) climbing fibers multi-innervate the Purkinje cell somas. At P24, tdTomato(+) climbing fibers mono-innervate Purkinje cell dendrites. Blue = calbindin (Purkinje cells); Purple = tdTomato; Green = vGluT2. **E.** Representative images from cKO mice. At P6, tdTomato(+) climbing fibers multi-innervate the Purkinje cell somas. At P24, tdTomato(+) climbing fibers mono-innervate Purkinje cell dendrites. Fewer tdTomato(+) climbing fibers are observed in P24 cKO cerebellums than in control cerebellums. Blue = calbindin (Purkinje cells); Purple = tdTomato; Green = vGluT2. **F.** No significant difference between groups at P6 (p=0.4902) or P9 (p=0.0713), but fewer tdTomato(+) climbing fibers in cKO cerebella compared to controls at P12 (p < 0.0001), P18 (p < 0.0001), and P24 (p = 0.0001). Within each genotype, the number of tdTomato(+) climbing fibers decreased over time relative to P6 (# indicates a significant difference from the same genotype at P6; Control: P9: p=0.0137, P12: p=0.0281, P18: p=0.0116, P24: p=0.0002; cKO: P9: p<0.0001, P12: p<0.0001, P18: p<0.0001, P24: p<0.0001). Control: P6 – N=5 mice, n=21 sections, *n*=9,171 calbindin(+) cells; P9 – N=4, n=29, *n*=16,038; P12 – N=3, n=22, *n=*10,584; P18 – N=5, n=20, *n=*7,683; P24 – N=4, n=21, *n=*9,376; cKO: P6 – N=3, n=18, *n*=8,273; P9 – N=5, n=34, *n*=9,376; P12 – N=3, n=18, *n=*9,491; P18 – N=3, n=18, *n=*7,621; P24 – N=3, n=18, *n=*8,452. **G.** Proportion of Purkinje cells with only tdTomato(+) climbing fibers at P6. No difference between genotypes. (p=0.2720, control: N=4, n=27, *n=*19,417; cKO: N=3, n=17, *n=*12,533). **H.** Quantification of tdTomato(+) climbing fibers in *Pdx1^+/+^;Rosa26^tdTomato^* mice (N=3, n=18, *n*=7,229). **I.** Quantification of tdTomato(+); vGluT2(+) climbing fibers are found in cKO cerebella (N=3, n=17, *n*=255). **J.** At P12, tdTomato(+), vGluT2(–) climbing fibers are innervating Purkinje cell dendrites, while vGluT2(+) climbing fibers are observed on soma. **K.** Quantification showed a reduction in tdTomato(+) climbing fiber innervation height in the cKO cerebella. (p = 0.0040; control: N=4, n=29, *n=*115; cKO: N=3, n=17, *n=*61). **L.** Quantification showed no reduction in vGluT2(+) climbing fiber innervation height in the cKO cerebella. (p = 0.5915; control: N=4, n=21, *n=*105; cKO: N=3, n=18, *n=*90) **M.** Schematic illustrates at P6, control and cKO cerebella have similar percentages of tdTomato(+) climbing fibers. At P24, there are fewer tdTomato(+) climbing fibers in the cKO cerebella.

Only mice carrying two floxed vGluT2 alleles are affected by this manipulation. Therefore *Pdx1^Cre^;Rosa26^tdTomato^* mice carrying either one floxed and one wild-type allele (*vGluT2^fl/+^*) or two wild-type alleles (*vGluT2^+/+^*) served as controls, whereas mice with two floxed copies (*vGluT2^fl/fl^*) are our experimental conditional knockout (cKO) animals in this experiment (Figure 2B).

We first confirmed that this strategy selectively eliminated vGluT2 expression from tdTomato(+) climbing fibers as early as P6. In control mice, tdTomato(+) fibers co-localized with vGluT2 labeling (Figure 2C, upper right Purkinje cell). In contrast, tdTomato(+) fibers in cKO mice lacked detectable vGluT2 expression (Figure 2C, lower right Purkinje cell). Thus, conditional deletion of vGluT2 specifically abolishes vGluT2(+)-dependent neurotransmission in the tdTomato(+) climbing fiber population, allowing us to assess how loss of neurotransmission affects competitive input selection.

We next characterized tdTomato(+) climbing fiber innervation of Purkinje cells across development in control mice at P6 (postnatal day 6), P9, P12, P18, and P24 (Supplemental figure 3). At P6, most Purkinje cells were multiply innervated by both tdTomato(+) and tdTomato(–) climbing fibers, and approximately 80% of Purkinje cells received at least one tdTomato(+) input (Figure 2D,F). By P9, multi-innervation was largely absent, reflecting completion of climbing fiber selection. Correspondingly, the proportion of Purkinje cells innervated by tdTomato(+) climbing fibers decreased to approximately 50% and remained stable through all later time points. We found no difference between mice carrying one or two wild-type copies of *vGluT2* (*vGluT2^fl/+^* versus *vGluT2^+/+^*) in the proportion of Purkinje cells innervated by tdTomato(+) climbing fibers (t-test: p = 0.6199; *vGluT2^fl/+^*: n = 9 mice, μ = 50.0%, σ = 13.1%; *vGluT2^+/+^*: n = 7 mice, μ = 53.0%, σ = 9.1%). These findings confirm that in control mice, tdTomato(+) climbing fibers normally undergo competitive selection and are retained as the sole climbing fiber input in approximately half of Purkinje cells.

We then examined climbing fiber selection in cKO mice lacking vGluT2 in tdTomato(+) climbing fibers. Elimination of vGluT2 did not disrupt global cerebellar morphology (Supplemental Figure 2). Similar to controls, most Purkinje cells at P6 were multiply innervated by both tdTomato(+) and tdTomato(–) climbing fibers, and approximately 80% received at least one tdTomato(+) input (Figure 2E,F). Thus, loss of vGluT2 did not affect the initial establishment of olivocerebellar connections.

However, unlike control mice at P9, many Purkinje cells in cKO mice remained multiply innervated by both tdTomato(+) and tdTomato(–) climbing fibers, indicating a delay in climbing fiber selection. Persistent multi-innervation was observed in four of five P9 cKO mice. In each case, a similar spatial pattern emerged: Purkinje cells in medial (vermal) regions were predominantly mono-innervated, whereas more lateral regions, including the paravermis and hemispheres, retained substantial multi-innervation (Supplemental Figure 4). This delayed refinement was resolved by P12, after which Purkinje cells in cKO mice were significantly less likely to select tdTomato(+) climbing fibers compared with controls.

Together, these results demonstrate that vGluT2-dependent neurotransmission is not required for the initial formation of olivocerebellar connections. However, it contributes to the timing and outcome of competitive climbing fiber selection, such that loss of neurotransmission delays input refinement and reduces the likelihood that affected climbing fibers are selected for mono-innervation.

### Climbing fibers lacking vGluT2 can be selected for mono-innervation

Although greatly reduced, tdTomato(+) climbing fibers were not completely excluded from selection in cKO mice. Some Purkinje cells still retained tdTomato(+) climbing fibers as their sole input. This could not be explained by a lack of competition from functional climbing fibers. At P6, before selection was complete, only 0–8% of Purkinje cells in either control or cKO mice were contacted exclusively by tdTomato(+) climbing fibers (Figure 2G). This proportion was far lower than the percentage of Purkinje cells mono-innervated by tdTomato(+), vGluT2(–) climbing fibers in P12-P24 cKO mice, indicating that these inputs were selected despite the presence of competing vGluT2-positive climbing fibers during early development.

We also ruled out technical explanations for the persistence of tdTomato(+) inputs. Ectopic, Cre-independent tdTomato expression was extremely rare, with fewer than 1% of climbing fibers labeled in mice lacking a Cre allele (Figure 2H). Likewise, incomplete recombination was minimal, as only ∼3% of tdTomato(+) climbing fibers had detectable vGluT2 expression in cKO mice (Figure 2I). These controls confirm that Cre-dependent labeling and manipulation were highly specific and that off-target effects are unlikely to account for the selection of tdTomato(+) climbing fibers.

Consistent with this interpretation, we occasionally observed multiply innervated Purkinje cells in P12 cKO mice. In some cases, a tdTomato(+), vGluT2(–) climbing fiber had already extended onto the Purkinje cell dendritic tree, whereas a competing tdTomato(–), vGluT2(+) climbing fiber remained restricted to the Purkinje cell soma (Figure 2J). This arrangement is characteristic of the tdTomato(+) climbing fiber being selected for long-term retention, suggesting that climbing fibers lacking vGluT2-mediated neurotransmission can still win the competition for Purkinje cell innervation. Because these examples were exceedingly rare, we could not quantify them further.

These findings indicate that vGluT2-dependent neurotransmission is not cell-intrinsically necessary but does contribute to the climbing fibers’ competitiveness during input selection. Next, we investigated whether climbing fibers’ likelihood of input selection is also compromised when most neighboring climbing fibers also lack vGluT2, *Ptf1a^Cre^;vGluT2^fl/fl^* mice.^18^ Similar to our observations in *Pdx1^Cre^;vGluT2^fl/fl^* mice (Supplemental Figure 2), loss of vGluT2-dependent neurotransmission from the *Ptf1a*-domain did not result in gross morphological abnormalities in the cerebellum.

We confirmed that *Ptf1a^Cre^* efficiently removed vGluT2 from most climbing fibers. In *Ptf1a^Cre^;vGluT2^fl/fl^* mice, only 8.1 ± 1.4% of Purkinje cells were innervated by vGluT2(+) climbing fibers (N = 4 mice, n = 12 sections). Because *Ptf1a^Cre^*is expressed in both climbing fibers and Purkinje cells,^29^ unlike *Pdx1^Cre^* (Figure 1), a Cre-dependent reporter could not be used to selectively trace climbing fibers. Instead, we used cocaine- and amphetamine-regulated transcript (CART), which labels a subset of climbing fibers predominantly located in lobule X.^30,31^ We quantified the proportion of Purkinje cells innervated by CART-positive climbing fibers in *Ptf1a^Cre^;vGluT2^fl/fl^* mice and *vGluT2^fl/fl^*controls (Supplemental Figure 5A,B). The proportion of CART-positive climbing fiber innervation was significantly lower in cKO mice than in controls (Supplemental Figure 5C).

These findings independently support our results in *Pdx1^Cre^;vGluT2^fl/fl^*mice. Even when only a small fraction of climbing fibers retain vGluT2 expression, climbing fibers lacking vGluT2-mediated neurotransmission are less likely to be selected during competitive input refinement. Together, these data demonstrate that neurotransmission is not strictly required for climbing fiber retention but strongly biases the outcome of competitive input selection.

### Neurotransmission promotes climbing fiber innervation height

A previous study suggested that neurotransmission regulates how far climbing fibers extend into the molecular layer.^32^ To test this, we quantified climbing fiber height relative to molecular layer thickness. In *Pdx1^Cre^;vGluT2^fl/fl^*cKO mice, tdTomato(+) climbing fibers extended approximately 10% less into the molecular layer than tdTomato(+) climbing fibers in control mice (Figure 2K), confirming that neurotransmission promotes climbing fiber dendritic innervation. In contrast, neighboring tdTomato(–), vGluT2(+) climbing fibers showed no difference in innervation height between control and cKO mice (Figure 2L), indicating that this effect is specific to climbing fibers lacking neurotransmission. To further test whether this effect is cell autonomous, we examined CART(+) climbing fibers in *Ptf1a^Cre^;vGluT2^fl/fl^* mice, in which most climbing fibers lack vGluT2. CART(+) climbing fibers in these mice also exhibited reduced innervation height compared with CART(+) climbing fibers in control littermates (Supplemental Figure 5D). Together, these findings suggest that vGluT2-mediated neurotransmission promotes the expansion of climbing fiber dendritic arbors within the molecular layer.

### Climbing fiber selection promotes inferior olive neuron survival

Many brain regions make supernumerous neurons in addition to supernumerous synaptic contacts, and it is widely assumed that only neurons that are used in active circuits survive ^33,34^ whereas other neurons do not.^35^ Programmed cell-death also occurs in the inferior olive,^36,37^ the source of climbing fiber projections to the cerebellum, but this process largely precedes climbing fiber input selection during normal development suggesting that natural inferior olive neuron death occurs through a different mechanism than input selection.^37^ Although inferior olive neuron survival depends on Purkinje cell survival,^38^ the presence of supernumerary climbing fiber inputs does not prevent developmental cell death.^37^ These observations raise the question of whether vGluT2-dependent input selection is required for inferior olive neuron survival.

To answer this question, we examined inferior olive neuron survival in mice lacking vGluT2 in tdTomato(+) neurons. At P6, the proportion of tdTomato(+) neurons was similar between control and cKO mice (Figure 3A-C), indicating that vGluT2-dependent neurotransmission is not required for inferior olive neuron survival prior to climbing fiber input selection.

**Figure 3.**
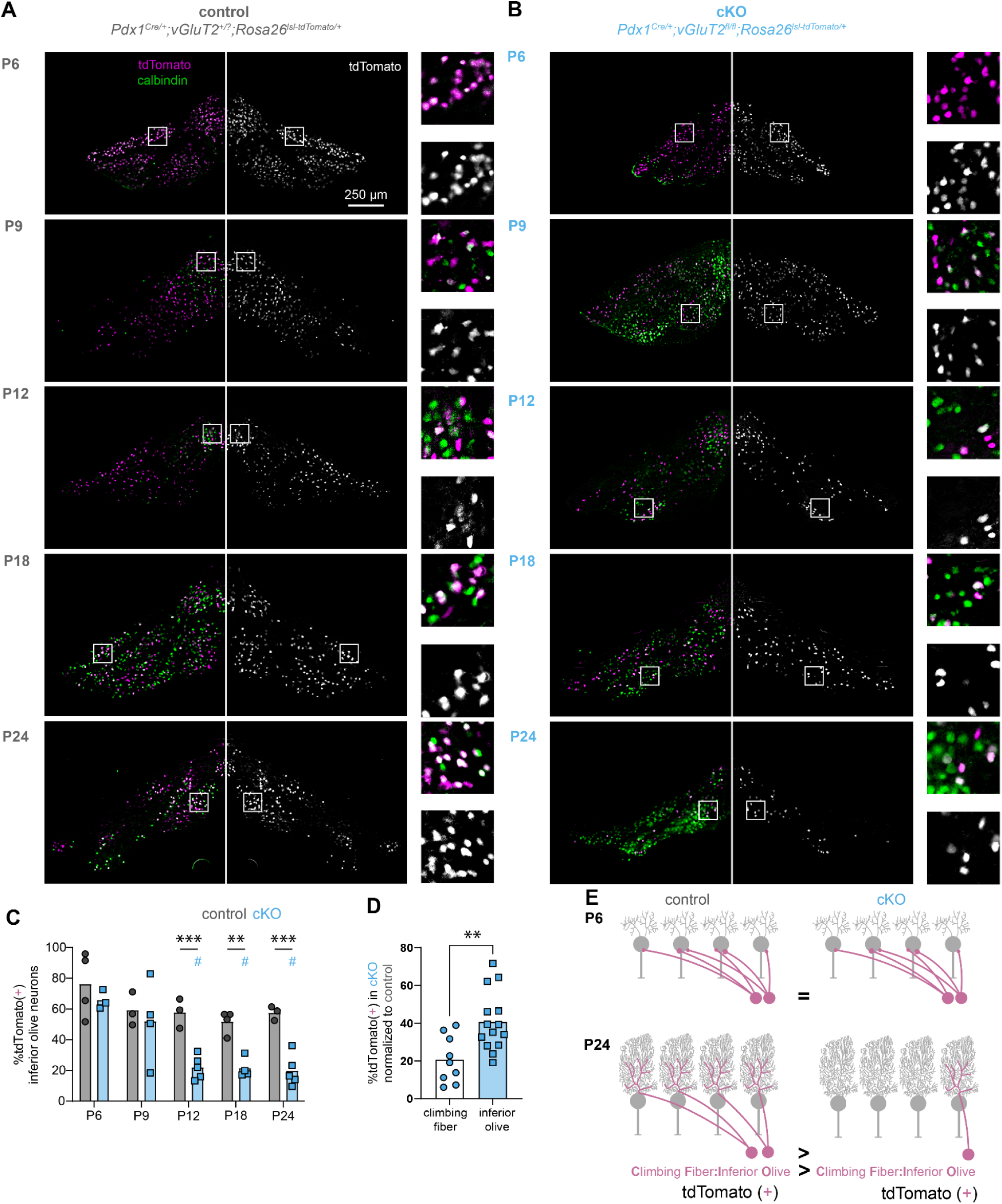
tdTomato labeling reveals reduced inferior olivary neuron survival in *Pdx1^Cre^;vGluT2^fl/fl^* mice. **A.** Representative images reveal robust tdTomato (purple) labeling of *Pdx1*(+) inferior olive nuclei and calbindin (green) labeling of *Pdx1*(-) inferior olive nuclei across all examined developmental stages. **B.** Representative images show abundant tdTomato and calbindin labeling at P6 and P9; however, the proportion number of tdTomato(+) neurons was diminished in the inferior olive at P12 and across all subsequent developmental time points examined. **C.** No significant difference between proportion of tdTomato(+) inferior olive neurons between control and cKO mice at P6 (p=0.3131) or P9 (p=0.4850), but fewer tdTomato(+) inferior olive neurons in cKO cerebella compared to controls at P12 (p = 0.0010), P18 (p =0.0032), and P24 (p = 0.0006). Only in the cKO mice, did the proportion of tdTomato(+) inferior olive neurons decrease over time relative to P6 (# indicates a significant difference from the same genotype at P6; P9: p=0.4454, P12: p=0.0004, P18: p=0.0006, P24: p=0.0002). Control: P6 – N=4 mice, n=20 sections, *n*=8,881 calbindin(+) cells; P9 – N=3, n=21, *n*=12,939; P12 – N=3, n=14, *n=*5,527; P18 – N=3, n=21, *n=*7,403; P24 – N=3, n=15, *n=*4,682; cKO: P6 – N=3, n=11, *n*=3,693; P9 – N=4, n=23, *n*=12,285; P12 – N=5, n=28, *n=*8,074; P18 – N=4, n=19, *n=*7,417; P24 – N=5, n=33, *n=*7,477. **D.** Relative proportion of tdTomato(+) climbing fibers and inferior olive neurons after climbing fiber selection has occurred (data from P12, P18, P24 cKO from panel 2F and 3C). Data points are normalized to average observed in control mice from the same cell-type and P12, P18, and P24. The relative proportion of selected tdTomato(+) climbing fiber is smaller than the relative proportion of surviving tdTomato(+) inferior olive neurons (p=0.0043). **E.** Schematic illustrating tdTomato(+) climbing fibers and inferior olive neurons in control and cKO cerebella. At P6, control and cKO mice exhibit similar proportions of tdTomato(+) climbing fibers and inferior olive neurons. At P24, proportion of tdTomato(+) climbing fibers and inferior olive neurons is reduced in cKO mice. However, because the relative reduction in tdTomato(+) climbing fiber is larger than the relative reduction in inferior olive neurons, it is likely that the proportion of climbing fibers per inferior olive neurons is also reduced in cKO mice.

We then analyzed P9 mice, a developmental stage at which some, but not all, Purkinje cells have completed climbing fiber selection (Supplemental Figure 4). At this age, the proportion of tdTomato(+) inferior olive neurons reflected the state of climbing fiber competition in the cerebellar cortex. Mice that still exhibited ongoing climbing fiber competition had proportions of tdTomato(+) neurons comparable to controls (Figure 3, Supplemental Figure 4), whereas the one *Pdx1^Cre^;vGluT2^fl/fl^*cKO mouse in which most Purkinje cells had already completed climbing fiber selection showed a marked reduction in tdTomato(+) neurons, approaching the levels observed at P12 and later ages (Figure 3C).

Importantly, even though we observed regional variations in Purkinje cell input selection in P9 cerebella, with medial regions selecting their climbing fibers prior to the more lateral regions, we did not observe any regional differences in density of tdTomato(+) neurons in corresponding regions in the inferior olive (Supplemental Figure 4). These findings suggest that the timing of climbing fiber selection and inferior olive neuron loss varies somewhat across animals, but that inferior olive neuron death does not occur until after Purkinje cells have selected their climbing fiber inputs.

To compare climbing fiber selection and inferior olive neuron survival after input selection was complete, we analyzed P12, P18, and P24 mice. For each time point, values from cKO mice were normalized to the mean value observed in age-matched controls. The relative proportion of selected tdTomato(+) climbing fibers was substantially lower than the relative proportion of surviving tdTomato(+) inferior olive neurons. This discrepancy indicates that the number of climbing fibers contributed by each surviving inferior olive neuron is reduced in cKO mice. Thus, loss of vGluT2-dependent neurotransmission appears to impair climbing fiber competitiveness, thereby reducing the parent neurons’ likelihood for survival.

Together, our findings suggest that vGluT2-dependent neurotransmission promotes the competitive success of olivocerebellar climbing fibers during input selection. Furthermore, the association between successful climbing fiber selection and increased inferior olive neuron survival raises the possibility that climbing fiber input selection contributes to the maintenance and survival of their parent neurons.

### Competing, unmanipulated climbing fibers expand their Purkinje cell innervation territory

If the reduced Purkinje cell innervation by tdTomato(+), vGluT2(–) climbing fibers reflects a loss of competitive advantage, then competing vGluT2(+) climbing fibers should expand their innervation territory. To test this hypothesis, we quantified the proportion of Purkinje cells innervated by vGluT2(+) climbing fibers at P12, when climbing fiber mono-innervation has been established, and in adulthood (>P56) to determine whether further territorial expansion occurs later in life.

In control mice (*vGluT2^fl/fl^* mice), vGluT2(+) climbing fiber innervation was dense and uniform at both ages examined (Figure 4A–B). In contrast, cKO mice (*Pdx1^Cre/+^;vGluT2^fl/fl^* mice) displayed a mosaic pattern of vGluT2(+) and vGluT2(–) climbing fiber territories at P12 (Figure 4C,E), which persisted into adulthood (Figure 4D,F). These observations indicate that not all Purkinje cells are innervated by vGluT2(+) climbing fibers in cKO mice.

**Figure 4.**
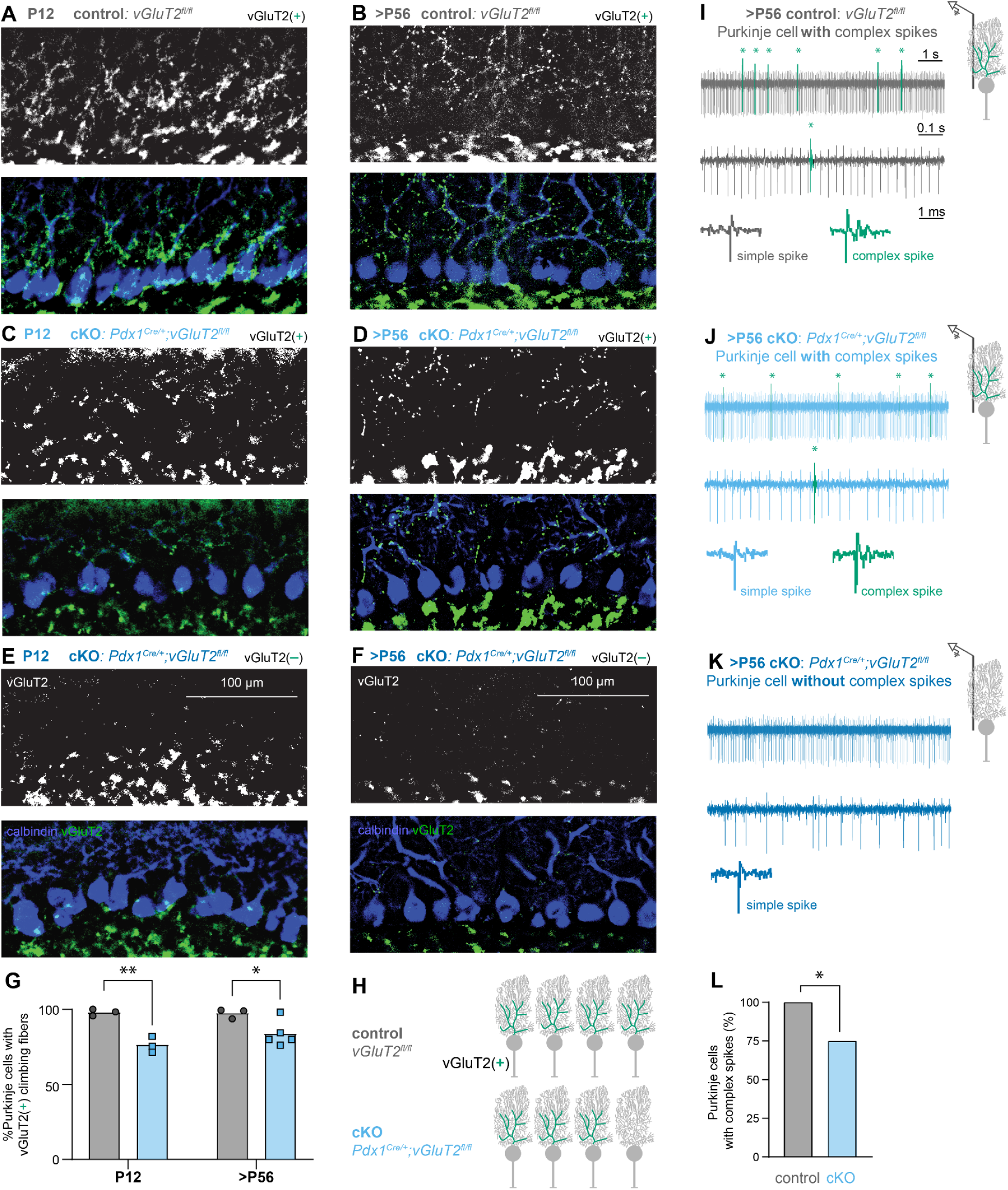
*Pdx1^Cre^;vGluT2^fl/fl^* cerebella exhibit fewer vGluT2(+) climbing fibers than controls, with expression remaining stable from early postnatal development to adulthood. **A, C, E.** Left side shows that at P12, the *Pdx1^Cre^;vGluT2^fl/fl^* cerebellum (cKO) contains regions with vGluT2(+) climbing fibers comparable to controls, alongside areas lacking vGluT2(+) climbing fibers. Blue = calbindin (Purkinje cells); Green = vGluT2. **C** and **E** are from the same cerebellar section. **B, D, F.** Right side shows that >P56, the cKO cerebellum contains regions with vGluT2(+) climbing fibers comparable to controls, alongside areas lacking vGluT2(+) climbing fibers. Blue = calbindin (Purkinje cells); Green = vGluT2. **D** and **F** are from the same cerebellar section. **G.** Quantification confirmed consistent reduction of vGluT2 signal in the *Pdx1^Cre^;vGluT2^fl/fl^*cerebellum at P12 (p = 0.0024) and >P56 (p = 0.0127). P12 Control: N=3 mice, n=9 sections; P12 *Pdx1^Cre/+^; vGluT2^fl/fl^;* N=3, n=9; >P56 Control: N=3, n=10; >P56 *Pdx1^Cre^;vGluT2^fl/fl^*: N=5, n=10. **H.** Schematic illustrates that there are fewer vGluT2(+) climbing fibers in the cKO cerebellum compared to control. **I.** Representative trace of a Purkinje cell recording in a control mouse (Gray lines = simple spikes; Green lines = complex spikes). **J.** Representative trace of a Purkinje cell recording in an area with vGLuT2(+) climbing fibers in a *Pdx1^Cre^;vGluT2^fl/fl^*mouse (Light blue lines = simple spikes; Green lines = complex spikes). **K.** Representative trace of a Purkinje cell recording in an area lacking vGluT2(+) climbing fibers in a *Pdx1^Cre^;vGluT2^fl/fl^* mouse (Dark blue lines = simple spikes). In **I, J,** vGluT2(+) climbing fibers evoked complex spikes (CS) in green. The timescales are the same for I, J, and K. L. *Pdx1^Cre^;vGluT2^fl/fl^*mice exhibited a significant reduction in complex spikes incidence (75%) compared to controls (100%), representing a 25% absolute decrease (Fisher’s exact test, p = 0.0127).

To quantify the extent of territory expansion, we measured the proportion of Purkinje cells innervated by vGluT2(+) climbing fibers in cKO mice. Approximately 75% of Purkinje cells received vGluT2(+) climbing fiber input, consistent with an expansion of vGluT2(+) territory beyond its expected baseline representation (Figure 4G–H). However, vGluT2(+) climbing fibers never occupied the entire Purkinje cell population at either age. This finding is consistent with our earlier observation that a subset of tdTomato(+), vGluT2(–) climbing fibers was selected for Purkinje cell mono-innervation. Furthermore, the proportion of Purkinje cells innervated by vGluT2(+) climbing fibers was similar at P12 and >P56, indicating that the expanded territory was established during developmental climbing fiber selection and remained stable thereafter.

Together, these findings demonstrate that vGluT2(+) climbing fibers expand their innervation territory by outcompeting a subset of vGluT2(–) climbing fibers and that this altered pattern of connectivity persists into adulthood. These results provide further evidence that vGluT2-dependent neurotransmission contributes to climbing fiber competitive success during input selection. Consequently, selective elimination of neurotransmission from a subset of climbing fibers generates an olivocerebellar circuit in which vGluT2-expressing climbing fibers innervate most, but not all, Purkinje cells (Figure 4H).

### Climbing fiber neurotransmission is necessary for complex spikes in some Purkinje cells without changing overall firing patterns

Next, we investigated whether eliminating vGluT2 from a subset of climbing fibers alters Purkinje cell firing activity. Purkinje cells generate two distinct types of action potentials: simple spikes, which occur spontaneously, and complex spikes, which are driven by glutamatergic climbing fiber input and are characterized by a large initial spike followed by smaller spikelets (Figure 4I).^39,40^ Previous work demonstrated that eliminating vGluT2 from most climbing fibers in *Ptf1a^Cre^;vGluT2^fl/fl^* mice largely abolishes complex spikes while simple spike firing patterns are unchanged in adulthood.^18^ Here, we used in vivo electrophysiology to determine whether deleting vGluT2 from only a subset of climbing fibers similarly affects complex spike occurrence and alters simple or complex spike firing properties.

Consistent with our anatomical findings that nearly all Purkinje cells in control mice are innervated by vGluT2(+) climbing fibers (Figure 4A,B,G), all recorded Purkinje cells exhibited complex spikes (25/25 cells; Figure 4I). In contrast, only 15 of 20 Purkinje cells recorded from *Pdx1^Cre/+^;vGluT2^fl/fl^*cKO mice exhibited complex spikes (Figure 4J,K). This proportion closely matches the percentage of Purkinje cells innervated by vGluT2(+) climbing fibers (∼75%), suggesting that vGluT2(+) climbing fibers retain the ability to evoke complex spikes, whereas vGluT2(–) climbing fibers do not (Figure 4L).

We next examined simple and complex spike firing properties in Purkinje cells that exhibited complex spikes. We detected no differences between control and cKO mice in simple spike or complex spike firing rate, pattern, or regularity (Supplemental Figure 6A–F). Together, these findings provide functional evidence that loss of vGluT2 selectively disrupts climbing fiber-mediated complex spike generation in a subset of Purkinje cells, while largely preserving the firing properties of the Purkinje cells.

### Intersectional genetics demonstrates lineage-specific overlap between vGluT2 and inferior olive marking *Pdx1^Cre^* and *Ptf1a^Cre^*lines

Having established that eliminating vGluT2 from a subset of climbing fibers leads to atypical olivocerebellar circuit assembly, we next sought to determine how these circuit changes influence behavior. As a first step, we defined the neuronal populations affected by our genetic manipulation.

To do so, we employed an intersectional genetic labellinh strategy in which tdTomato expression is activated only after excision of two stop cassettes by FlpO, driven by the *vGluT2* promoter, and Cre, driven by the *Pdx1* promoter (Figure 5A). This approach labels only neurons with a history of both *Pdx1* and *vGluT2* expression (Figure 5A–B), thereby marking the intersectional domain affected in *Pdx1^Cre/+^;vGluT2^fl/fl^*conditional knockout mice.

**Figure 5.**
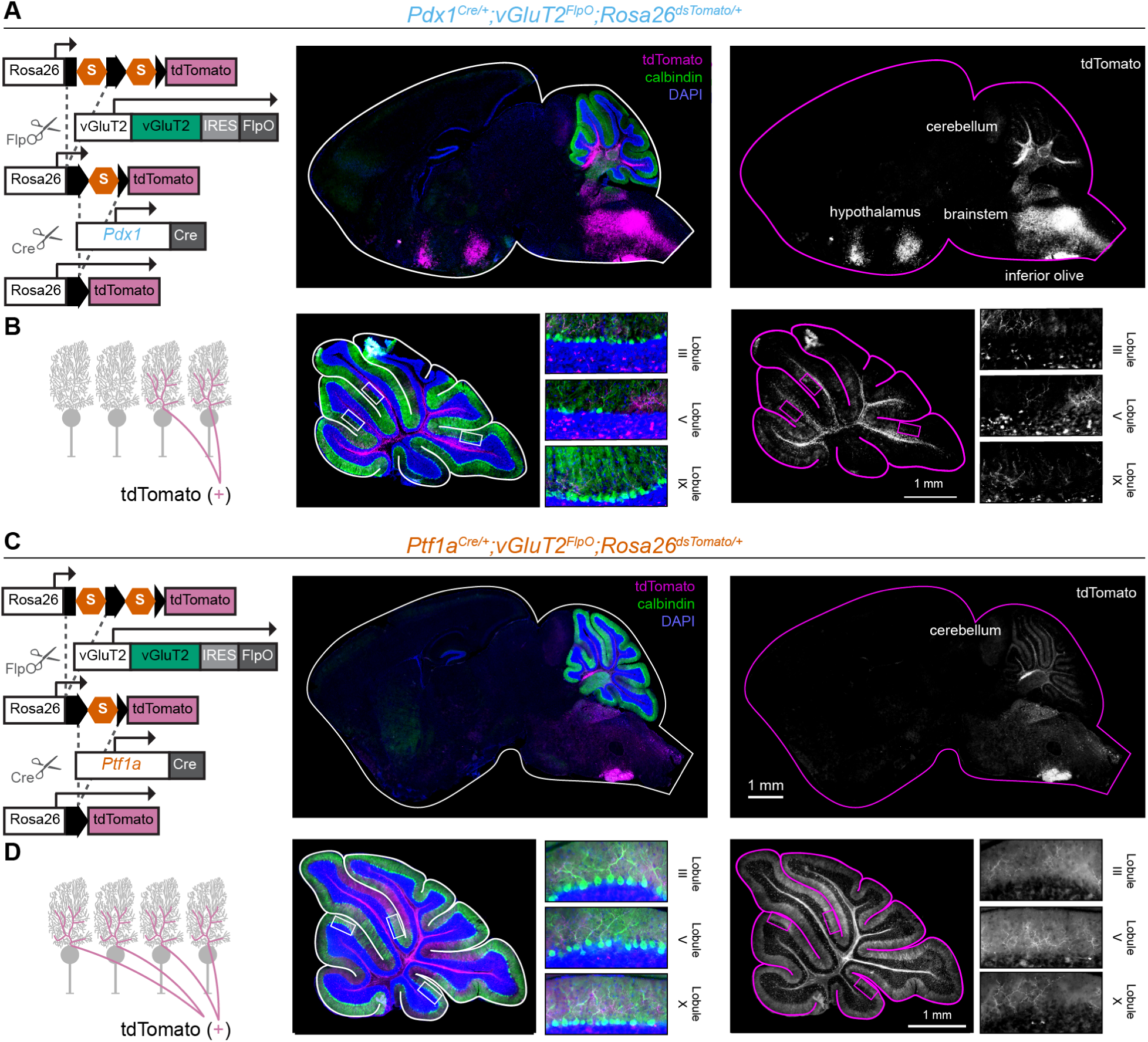
Intersectional genetic strategy for selective labeling of *Pdx1^Cre^*(+);*Vglut2*(+) and *Ptf1a ^Cre^* (+)*; Vglut2*(+) neurons with tdTomato reporter. **A.** Genetic strategy that expresses the tdTomato reporter in all neurons upon *Pdx1^Cre^ and Vglut2^FlpO^-*mediated recombinase. Right panels show sagittal sections of the brain where tdTomato expression is present in the inferior olive, brainstem, hypothalamus, and the cerebellar cortex. Blue = DAPI (cell nuclei); Green = calbindin (Purkinje cells); Purple = tdTomato (also in white, right). **B.** Schematic illustrates tdTomato expression in *Pdx1^Cre^;Vglut2^FlpO^* climbing fibers. Right panels show sagittal cerebellar sections where tdTomato expression is present throughout the cerebellar cortex but not all climbing fibers. Blue = DAPI (cell nuclei); Green = calbindin (Purkinje cells); Purple = tdTomato (also in white, right). **C.** Genetic strategy that expresses the tdTomato reporter in all neurons upon *Ptf1a^Cre^ and Vglut2^FlpO^-*mediated recombinase. Right panels show sagittal sections of the brain where tdTomato expression is present in the inferior olive and cerebellar cortex. Blue = DAPI (cell nuclei); Green = calbindin (Purkinje cells); Purple = tdTomato (also in white, right). **D.** Schematic illustrates tdTomato expression in *Ptf1a^Cre^;Vglut2^FlpO^* climbing fibers. Right panels show sagittal cerebellar sections where tdTomato expression is present throughout the cerebellar cortex in all climbing fibers. Blue = DAPI (cell nuclei); Green = calbindin (Purkinje cells); Purple = tdTomato (also in white, right top/bottom row). Images are representative for N = 3 mice.

Using this strategy, we identified tdTomato(+) neurons in the inferior olive, as well as a subset of tdTomato(+) climbing fibers and mossy fibers throughout the cerebellar cortex (Figure 5A–B). However, tdTomato(+) neurons were also present in several other brainstem nuclei and hypothalamic regions. These findings indicate that the *Pdx1^Cre/+^;vGluT2^fl/fl^* manipulation affects neural populations beyond the olivocerebellar circuit, raising the possibility that behavioral phenotypes in these mice could reflect alterations in non-cerebellar circuits.

To address this potential confound, we mapped the intersectional domain between vGluT2 and a second Cre driver, *Ptf1a^Cre^*, which is widely used to target inferior olive neurons (Figure 5C). Previous studies demonstrated that *Ptf1a^Cre^*labels nearly all olivocerebellar climbing fibers,^18^ consistent with our observation that only 8.1 ± 1.4% of Purkinje cells are innervated by vGluT2-positive climbing fibers in *Ptf1a^Cre/+^;vGluT2^fl/fl^* mice (Figure 5D). In contrast to the broader Pdx1 intersectional domain, overlap between *Ptf1a^Cre^*and *vGluT2* expression was restricted to inferior olive neurons and their climbing fiber projections within the cerebellar cortex (Figure 5C–D).

Given the specificity of the *Ptf1a^Cre/+^;vGluT2* intersectional domain, we used *Ptf1a^Cre/+^;vGluT2^fl/fl^* mice as a positive control for behavioral assays to determine whether task performance depends on vGluT2-mediated neurotransmission from climbing fibers.

### Atypical olivocerebellar circuits result in mild deficits in motor control

Having defined the neuronal populations affected by our conditional knockout models (Figure 5), we next investigated how atypical climbing fiber development influences motor behavior. We compared three groups of mice: control littermates, in which nearly all Purkinje cells are innervated by vGluT2-positive climbing fibers (97.5%); *Pdx1^Cre/+^;vGluT2^fl/fl^* mice, which exhibit partial innervation by vGluT2-positive climbing fibers (78.5%); and *Ptf1a^Cre^;vGluT2^fl/fl^* mice, in which very few Purkinje cells retain vGluT2-positive climbing fiber input (8.1%; Figure 6A). These models allowed us to determine whether the adaptive changes in circuit assembly observed in *Pdx1^Cre/+^;vGluT2^fl/fl^* mice are sufficient to preserve climbing-fiber-dependent motor function.

**Figure 6.**
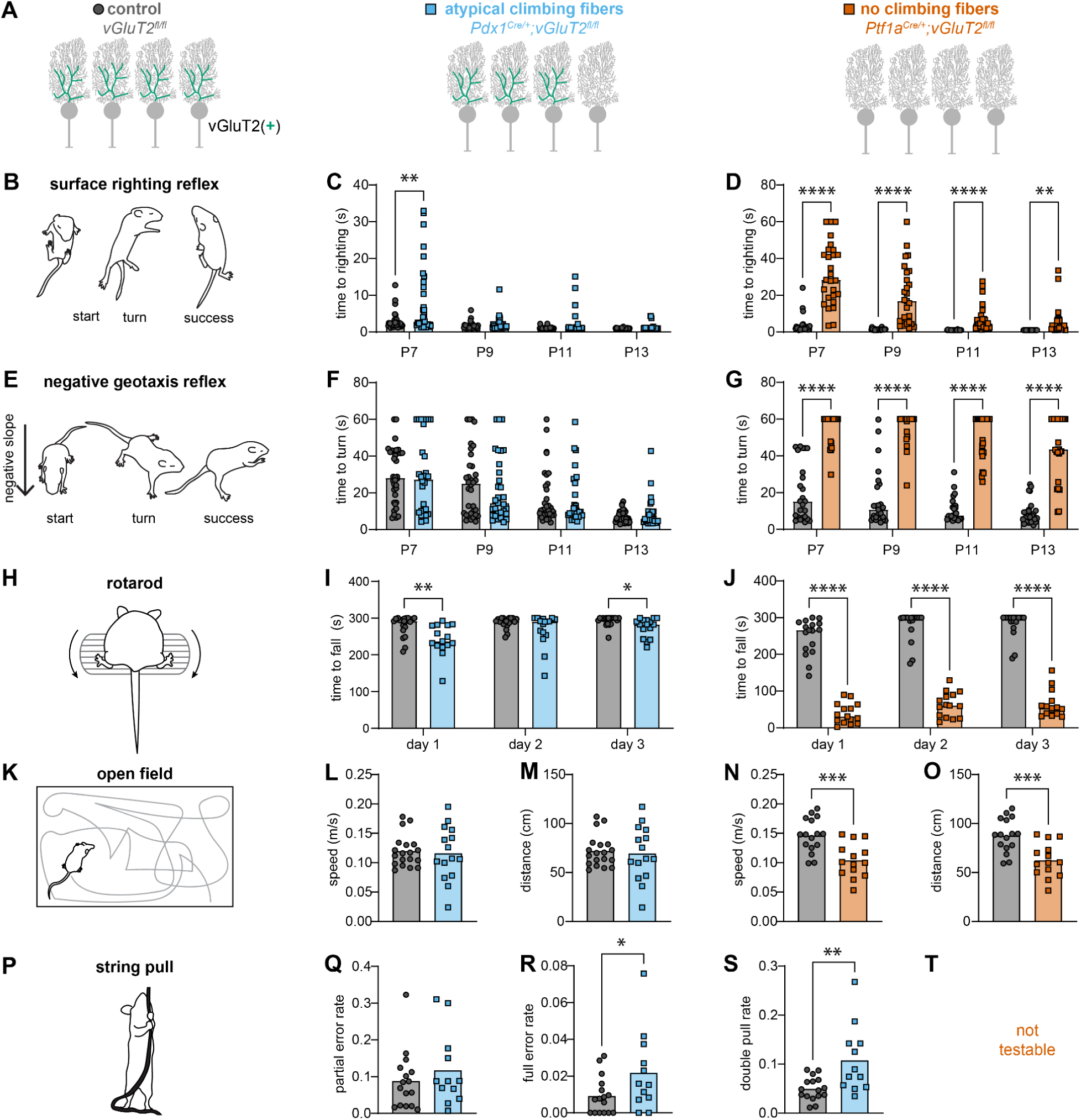
*Ptf1a^Cre^;vGluT2^fl/fl^* mice, but not *Pdx1^Cre^;vGluT2^fl/fl^* mice, exhibit severe motor dysfunction from early postnatal stages through adulthood. **A.** Schematic illustrates atypical vGluT2(+) climbing fiber development in the *Pdx1^Cre^;vGluT2^fl/fl^* cerebellum compared to control, and no vGluT2(+) climbing fibers in the *Ptf1a^Cre^;vGluT2^fl/fl^* cerebellum. **B.** Schematic illustration of surface righting reflex: the time to right themselves onto their four paws was measured. C. *Pdx1^Cre^;vGluT2^fl/fl^*mice exhibited slower motor reflexes to right onto all four paws compared to control littermates at P7 (p = 0.0032), but not at P9 (p = 0.2498), P11 (p = 0.0773), or P13 (p = 0.0750). D. *Ptf1a^Cre^;vGluT2^fl/fl^* mice exhibited slower motor reflexes to right onto their four paws compared to control littermates at P7 (p < 0.0001), P9 (p < 0.0001), P11(p < 0.0001), and P13 (p = 0.0039). **E.** Schematic illustrates the time to turn upward on a negative slope was measured. **F.** No differences were observed in the negative geotaxis reflex between the *Pdx1^Cre/+^; Vglut2^fl/fl^* and control littermates at P7 (p = 0.9892), P9 (p = 0.4684), P11(p = 0.8107), and P13(p = 0.2361). G. *Ptf1a^Cre^;vGluT2^fl/fl^* mice exhibited slower motor reflexes to turn upwards compared to control littermates at P7 (p < 0.0001), P9 (p < 0.0001), P11(p < 0.0001), and P13 (p < 0.0001). For **C,F**: Control: N = 34 mice (22M/12F); *Pdx1^Cre^;vGluT2^fl/fl^*: N = 31 mice (14M/17F); For **D,G** Control: N = 25 mice (17M/8F); *Ptf1a^Cre^;vGluT2^fl/fl^*: N = 27 mice (9M/18F). **H.** Schematic illustrates the duration of maintaining locomotion in response to increasing speed. I. *Pdx1^Cre^;vGluT2^fl/fl^*mice fell off the accelerating ramp faster than their control littermates on Day 01 (p = 0.0077) and Day 03 (p = 0.0259), but not Day 02 (p = 0.1189). J. *Ptf1a^Cre^;vGluT2^fl/fl^* mice fell off the accelerating ramp faster than their control littermates on Day 01 (p < 0.0001), Day 02 (p < 0.0001), and Day 03 (p < 0.0001). **K.** Schematic illustrates the distance and speed of travel in an open apparatus was measured. **L.** No differences were observed in the speed of travel between the *Pdx1^Cre^;vGluT2^fl/fl^* and control littermates (p = 0.7735). **M.** No differences were observed in the distance covered between the *Pdx1^Cre^;vGluT2^fl/fl^* and control littermates (p = 0.7578) For **L,M**: Control: N = 19 mice (8M/11F), *Pdx1^Cre^;vGluT2^fl/fl^*: N = 15 mice (8M/7F). N. *Ptf1a^Cre^;vGluT2^fl/fl^* mice traveled at slower speeds compared to control littermates (p = 0.0005). O. *Ptf1a^Cre^;vGluT2^fl/fl^* mice covered less distance compared to control littermates (p = 0.0005). For **N,O**: Control: N = 16 mice (8M/8F), *Ptf1a^Cre^;vGluT2^fl/fl^*: N = 14 mice (6M/8F). **P.** Schematic representation of a mouse performing the string-pull test to assess skilled forelimb motor coordination. **Q.** Quantification of partial miss rate revealed no significant differences between *Pdx1^Cre^;vGluT2^fl/fl^*and control littermates (p = 0.3909). **R.** Quantification of full miss rate revealed more eros in *Pdx1^Cre^;vGluT2^fl/fl^*than control littermates (p = 0.0493). **S.** Quantification of the double miss rate showed elevated error rates in *Pdx1^Cre^;vGluT2^fl/fl^*mice relative to control littermates (p = 0.0038). For **Q-S:** Control: N = 16 mice (12M/4F), *Pdx1^Cre^;vGluT2^fl/fl^*: N = 12 mice (8M/4F)). **T.** String pull assay could not be performed in the *Ptf1a^Cre/+^; Vglut2^fl/fl^* mice. Grey circles represent data points from control mice; Blue squares represent data points from *Pdx1^Cre^;vGluT2^fl/fl^* mice; Orange squares represent data points from *Ptf1a^Cre/+^; Vglut2^fl/fl^* mice.

To assess motor performance during development, we examined two postnatally acquired motor reflexes known to depend on cerebellar function.^41,42^ First, we measured the surface righting reflex by placing pups on their backs and recording the time required to return to all four paws (Figure 6B). *Pdx1^Cre/+^;vGluT2^fl/fl^* pups exhibited only a mild delay only at P7 compared to control littermates (Figure 6C),, with no differences detected at P9, P11, or P13. In contrast, *Ptf1a^Cre^;vGluT2^fl/fl^*pups showed severe impairments and required significantly longer to turn at every age examined (Figure 6D).

We next assessed the negative geotaxis reflex by placing pups head-down on an inclined surface and measuring the latency to rotate upward (Figure 6E). *Pdx1^Cre/+^;vGluT2^fl/fl^* pups performed similarly to controls at all ages tested (Figure 6F). By contrast, *Ptf1a^Cre^;vGluT2^fl/fl^* pups exhibited pronounced deficits, requiring significantly more time to orient upward (Figure 6G). Together, these findings indicate that atypical climbing fiber development causes only mild impairments in cerebellar-dependent motor reflexes during the early postnatal period.

We then asked whether motor performance remained affected in adulthood. To evaluate motor coordination and balance, mice were tested on an accelerating rotarod, and latency to fall was measured (Figure 6H). *Pdx1^Cre/+^;vGluT2^fl/f^* mice displayed mild deficits, falling slightly earlier than controls on testing days 1 and 3 (Figure 6I). In contrast, *Ptf1a^Cre^;vGluT2^fl/fl^* mice exhibited severe motor impairments, falling significantly sooner than control littermates on all testing days (Figure 6J).

To assess general locomotor activity, we performed an open-field assay and measured total distance traveled and movement speed (Figure 6K). *Pdx1^Cre/+^;vGluT2^fl/fl^* mice did not differ significantly from controls on either measure (Figure 6L,M). However, *Ptf1a^Cre^;vGluT2^fl/fl^*mice traveled shorter distances and moved more slowly than control littermates (Figure 6N,O). Together, these results demonstrate that while climbing fiber neurotransmission is critical for normal motor behavior, the adaptive circuit reorganization observed in *Pdx1^Cre/+^;vGluT2^fl/fl^* mice largely preserves gross motor function.

Finally, we investigated whether more subtle deficits emerged during a fine motor task. In the string-pulling assay, mice stand on their hindlimbs and use coordinated forelimb movements to pull a suspended string downward (Figure 6P). *Pdx1^Cre/+^;vGluT2^fl/fl^* mice were able to perform this task, whereas *Ptf1a^Cre^;vGluT2^fl/fl^* mice lacked the balance and coordination required for successful testing (Figure 6T).

Although *Pdx1^Cre/+^;vGluT2^fl/fl^* mice completed the task, they exhibited measurable deficits compared with control littermates. They did not show an increase in partial misses during an attempted grasp (Figure 6Q). However, they completely missed the string more frequently (Figure 6R) and, perhaps to compensate, more often adopted a bilateral pulling strategy, using both forepaws simultaneously rather than the alternating paw-over-paw movements typically used by control mice (Figure 6S). Because the paw-over-paw strategy is more efficient for advancing the string, these findings suggest that *Pdx1^Cre/+^;vGluT2^fl/fl^* mice compensate for impaired fine motor coordination by relying on a less efficient movement strategy.

Together, these results indicate that adaptive climbing fiber circuit assembly largely preserves gross motor behaviors but does not fully restore fine motor control, resulting in subtle deficits in movement precision and motor strategy.

### Atypical olivocerebellar circuits do not impair social behaviors

To further define the behavioral consequences of altered climbing fiber development, we investigated the contribution of climbing fibers to social behaviors. Using the same *Pdx1^Cre/+^;vGluT2^fl/fl^*and *Ptf1a^Cre^;vGluT2^fl/fl^* mouse models examined in our motor studies, we assessed social behaviors from early postnatal development through adulthood (Figure 7A).

**Figure 7.**
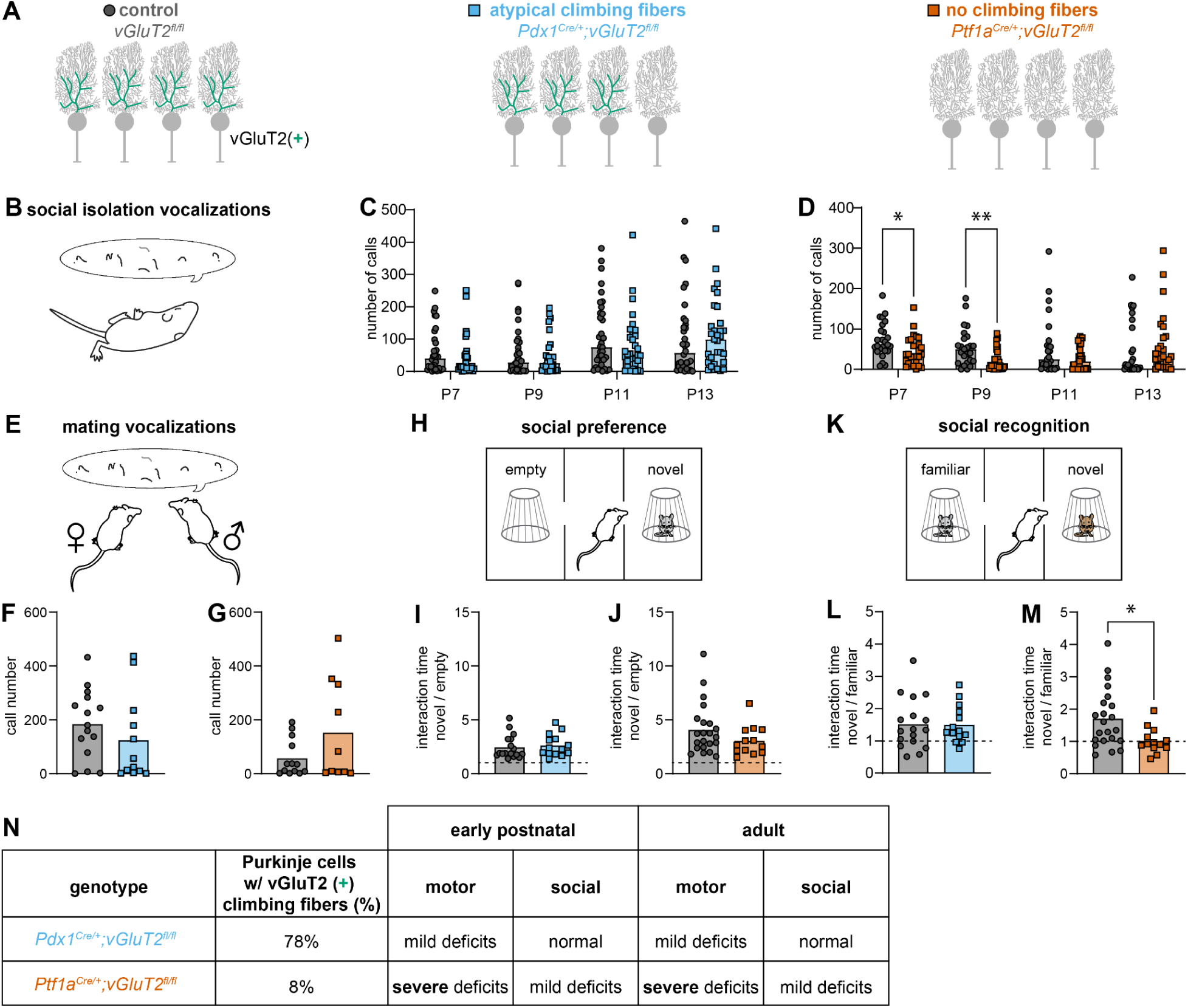
*Ptf1a^Cre^;vGluT2^fl/fl^* mice, but not *Pdx1^Cre^;vGluT2^fl/fl^* mice, exhibit deficits in social communication during early postnatal development and impaired social recognition in adulthood. **A.** Schematic illustrates atypical vGluT2(+) climbing fiber development in the *Pdx1^Cre^;vGluT2^fl/fl^* cerebellum compared to control, and no vGluT2(+) climbing fibers in the *Ptf1a^Cre^;vGluT2^fl/fl^*cerebellum. **B.** Schematic illustrates the number of vocalization calls produced by pups after being separated from their nest. **C.** No differences were observed in the number of calls produced between the *Pdx1^Cre^;vGluT2^fl/fl^* and control littermates at P7 (p = 0.2934), P9 (p = 0.4676), P11 (p = 0.2293), and P13 (p = 0.6615). Control: N = 34 mice (22M/12F); *Pdx1^Cre^;vGluT2^fl/fl^*: N = 31 mice (14M/17F). D. *Ptf1a^Cre^;vGluT2^fl/fl^*mice exhibited a reduced number of calls compared to their control littermates at P7 (p = 0.0154) and P9 (p = 0.0044), but no differences were found in P11 (p = 0.1040) and P13 (p = 0.5670). Control: N = 25 mice (17M/8F); *Ptf1a^Cre^;vGluT2^fl/fl^*: N = 27 mice (9M/18F). **E.** Schematic illustrates the number of vocalization calls produced by adult male mice during courtship with an adult female mouse. **F.** No differences were found in the number of calls produced between the *Pdx1^Cre^;vGluT2^fl/fl^* and control littermates (p = 0.2979, Control: N = 15 mice (15M), *Pdx1^Cre^;vGluT2^fl/fl^*: N = 12 mice (12M). **G.** No differences were found in the number of calls produced between the *Ptf1a^Cre^;vGluT2^fl/fl^* and control littermates (p = 0.1034, Control: N = 13 mice (13M); *Ptf1a^Cre^;vGluT2^fl/fl^*: N = 10 mice (10M). **H.** Schematic illustrates a novel mouse placed on one side, and an empty cage placed on the other side of the 3-chamber, and the time spent socially interacting with either stimulus was measured. **I.** No differences were observed in the ratio of interaction time spent with the novel mouse compared to the empty cage between the *Pdx1^Cre^;vGluT2^fl/fl^* and control littermates (p = 0.6478, Control: N = 17 mice (7M/ 10F); *Pdx1^Cre^;vGluT2^fl/fl^*: N = 14 mice (8M/6F). **J.** No differences were observed in the ratio of interaction time spent with the novel mouse compared to the empty cage between the *Ptf1a^Cre^;vGluT2^fl/f^* mice their control littermates (p = 0.1601, Control: N = 22 mice (12M/10F); *Ptf1a^Cre^;vGluT2^fl/fl^*: N = 13 mice (4M/9F). **K.** Schematic illustrates the previous novel mouse becomes the familiar mouse, and a new novel mouse replaces the empty cage on the other side of the 3-chamber, and the time spent socially interacting with either stimulus was measured. **L.** No differences were observed in the ratio of interaction time spent with the novel mouse compared to the familiar mouse between the *Pdx1^Cre^;vGluT2^fl/fl^*and control mice (p = 0.9618, N = 17 mice (7M/10F); *Pdx1^Cre^;vGluT2^fl/fl^*: N = 14 mice (8M/6F). M. *Ptf1a^Cre^;vGluT2^fl/fl^* mice showed a lower preference ratio for novel mice over the familiar mouse compared to their control littermates (p = 0.0152, Control: N = 22 mice (12M/10F), t-test; *Ptf1a^Cre^;vGluT2^fl/fl^*: N = 13 mice (4M/9F). **N.** Summary of results. Grey circles represent data points from control mice; Blue squares represent data points from *Pdx1^Cre^;vGluT2^fl/fl^* mice; Orange squares represent data points from *Ptf1a^Cre^;vGluT2^fl/fl^* mice.

During early postnatal development, we measured ultrasonic vocalizations produced by pups after temporary separation from the nest, a behavior that serves as an instinctive call to the dam and has previously been shown to depend on climbing fiber signaling (Figure 7B).^41,43^ *Pdx1^Cre/+^;vGluT2^fl/fl^* pups pups did not differ from control littermates at any age examined (Figure 7C). In contrast, *Ptf1a^Cre^;vGluT2^fl/fl^* pups produced fewer vocalizations at P7 and P9, although these deficits were no longer apparent at P11 or P13 (Figure 7D). These findings indicate that partial disruption of climbing fiber development does not impair early-life social vocalizations.

We next examined vocalization behavior in adulthood using a courtship paradigm. Because male mice produce ultrasonic vocalizations during interactions with females,^43–45^ we measured vocalizations emitted by adult males in the presence of a female mouse (Figure 7E). Although variability was high within both knockout groups, neither *Pdx1^Cre/+^;vGluT2^fl/fl^* nor *Ptf1a^Cre^;vGluT2^fl/fl^* mice differed significantly from their control littermates in the number of vocalizations produced (Figure 7F,G). These findings suggest that climbing fiber signaling is not required for adult male courtship vocalizations.

Finally, we assessed social preference and social recognition using the three-chamber assay. During the first phase, mice were given a choice between investigating a novel mouse enclosed in a barred cup or an identical empty cup (Figure 7H). We quantified the proportion of time spent interacting with each stimulus. Neither *Pdx1^Cre/+^;vGluT2^fl/fl^* nor *Ptf1a^Cre^;vGluT2^fl/fl^*mice differed from controls in their preference for the social stimulus over the empty cup (Figure 7I,J), indicating intact social preference.

In the second phase, the previously novel mouse became the familiar stimulus and a new unfamiliar mouse was introduced (Figure 7K). As expected, control mice preferentially investigated the newly introduced mouse. *Pdx1^Cre/+^;vGluT2^fl/fl^* mice exhibited a similar preference, indicating intact social recognition memory. In contrast, *Ptf1a^Cre^;vGluT2^fl/fl^* mice failed to discriminate between the familiar and novel mice, suggesting impaired social recognition memory (Figure 7L). These findings support a role for climbing fibers in social recognition while indicating that the adaptive circuit reorganization observed in *Pdx1^Cre/+^;vGluT2^fl/fl^* mice is sufficient to preserve this behavior.

Taken together, the behavioral findings reveal a graded relationship between the extent of climbing fiber disruption and behavioral outcome (Figure 7N). *Pdx1^Cre/+^;vGluT2^fl/fl^* mice, in which approximately 78% of Purkinje cells retain vGluT2(+) climbing fiber input, exhibited only mild motor deficits from early postnatal development through adulthood and showed no detectable impairments in social behavior. In contrast, *Ptf1a^Cre^;vGluT2^fl/fl^*mice, in which only approximately 8% of Purkinje cells receive vGluT2(+) climbing fiber input, displayed severe motor impairments beginning in early postnatal life that persisted into adulthood, along with deficits in developmental vocalizations and adult social recognition memory. Overall, these findings demonstrate that climbing fibers contribute to both motor and social behavioral development and that behavioral outcomes are only minimal upon atypical climbing fiber assembly.

## DISCUSSION

In this study, we used conditional and intersectional genetic approaches to investigate how competitive climbing fiber input selection shapes circuit assembly and behavioral resilience. We found that neurotransmission is a key determinant of climbing fiber competitiveness during Purkinje cell input selection. Climbing fibers lacking vGluT2-dependent neurotransmission innervated fewer Purkinje cells and their parent neurons were less likely to survive, while neighboring unperturbed climbing fibers expanded their innervation territory. These findings demonstrate that neurotransmission-dependent competition influences both synaptic connectivity and neuronal survival during olivocerebellar circuit development.

Importantly, we show that this competitive developmental process promotes the formation of resilient cerebellar circuits. Despite substantial alterations in climbing fiber connectivity, mice retained most climbing-fiber-dependent motor and social behaviors, indicating that adaptive circuit reorganization can compensate for developmental perturbations. Together, our findings reveal that competitive input selection is not only a mechanism for refining neural connectivity but also a developmental strategy that enhances circuit robustness and supports the emergence of complex behaviors.

### Climbing fiber neurotransmission is not dispensable for competitive input selection

Climbing fiber input elimination and selection has long been proposed to be a competitive process in which synaptic strength and neurotransmission influence whether a climbing fiber is ultimately selected as the sole input to a Purkinje cell.^9^ Neuroanatomical and electrophysiological studies support this model, showing that smaller, weaker synapses are preferentially eliminated, whereas larger, stronger synapses are retained during development.^11,46^ Our findings provide direct experimental evidence that neurotransmission contributes to this competitive process. Eliminating vGluT2-dependent neurotransmission from a subset of climbing fibers reduced their competitive success, resulting in fewer innervated Purkinje cells (Figure 2) and allowing neighboring, unperturbed climbing fibers to expand their innervation territory (Figure 4). Moreover, even in the absence of substantial competition, as in mice lacking vGluT2 in most climbing fibers (Supplemental Figure 5), climbing fibers were less likely to establish Purkinje cell innervation. Together, these findings support the hypothesis that neurotransmission contributes to climbing fiber selection during the establishment of Purkinje cell mono-innervation.

Our conclusions differ from those in a recent study by Kao and colleagues, which proposed that neurotransmission is dispensable for climbing fiber input selection.^32^ These contrasting results may arise from important differences in experimental design. Our genetic approach produced a mosaic pattern of vGluT2 deletion throughout the cerebellar cortex and inferior olive while avoiding invasive manipulations that could perturb surrounding tissue or introduce regional biases. In addition, the combination of tdTomato reporter expression and vGluT2 immunolabeling enabled unambiguous identification of genetically perturbed neurons and their projections. This approach therefore provided a highly tractable model for studying competitive climbing fiber selection under physiological conditions.

A second distinction lies in how climbing fiber selection was quantified. Our analyses were based on unbiased assessments of climbing fiber innervation across more than 100,000 Purkinje cells, independent of climbing fiber morphology or anatomical location. In contrast, the viral strategy used by Kao and colleagues restricted analyses to a small subset of Purkinje cells that remained co-innervated by competing climbing fibers at P12.^32^ Motivated by their observations, we carefully examined our own P12 material and similarly identified rare examples of multi-innervated Purkinje cells (Figure 2J). Consistent with their findings, we observed occasional Purkinje cells in which a vGluT2(–) climbing fiber was selected for mono-innervation despite competition from a vGluT2(+) climbing fiber. These observations demonstrate that neurotransmission is not absolutely required for a climbing fiber to be selected. However, such events do not alter the overall conclusion that loss of neurotransmission substantially reduces competitive success. Because multi-innervated Purkinje cells were so uncommon at this stage, our dataset lacked sufficient examples to rigorously investigate the mechanisms underlying these exceptional cases. Future studies should determine how climbing fibers lacking neurotransmission can occasionally outcompete neighboring, unperturbed inputs.

Despite our diverging conclusions on the importance of neurotransmission for climbing fiber input selection, we independently replicate a key observation from Kao and colleagues: the extension of climbing fibers into the Purkinje cell dendritic arbor is strongly dependent on neurotransmission (Figure 2K; Supplemental Figure 5D).^32^ This agreement suggests that the effect of neurotransmission on climbing fiber growth and dendritic territory expansion is robust across experimental approaches.

One alternative explanation for the expansion of vGluT2(+) climbing fiber territory is that it occurs passively after the loss of neighboring vGluT2(–) inferior olive neurons. Our data instead support an active competitive mechanism. Climbing fiber selection was markedly more sensitive to vGluT2 deletion than inferior olive neuron survival, and alterations in climbing fiber connectivity preceded the loss of parent inferior olive neurons (Figures 2, 3, and Supplemental Figure 4). These observations suggest that successful climbing fiber selection promotes inferior olive neuron survival, rather than surviving climbing fibers expanding only after neighboring cells were lost due to cell death of the parental neuron.

Taken together, our findings demonstrate that vGluT2-dependent neurotransmission enhances the competitive success of climbing fibers during developmental input selection and contributes indirectly to the survival of their parent inferior olive neurons. These results challenge the conclusion that neurotransmission is dispensable for climbing fiber selection and instead support the longstanding hypothesis that neurotransmission and synaptic strength are important determinants of competitive olivocerebellar circuit assembly.

### Climbing fibers contribute to various motor and social behaviors

Numerous studies have established that climbing fiber signaling is essential for motor coordination and motor learning,^15,18,47^ yet its role in social behaviors has remained largely unexplored. The cerebellum receives two primary sources of glutamatergic input: direct climbing fiber synapses onto Purkinje cells and indirect mossy fiber input, which influences Purkinje cell activity via the granule cell–parallel fiber pathway.^48^ A recent study demonstrated that the mossy fiber–granule cell pathway is dispensable for several Purkinje cell-dependent cognitive behaviors, including the social preference task used in Figure 7H–J.^49^ Together with our findings, this suggests that social behaviors assessed in this paradigm depend more strongly on climbing fiber neurotransmission, thereby extending the known behavioral functions of climbing fiber–Purkinje cell signaling beyond the motor domain.

In motor systems, climbing fibers are thought to convey error or teaching signals critical for motor learning.^47,50,51^ Accordingly, we expected that disrupting climbing fiber signaling would primarily affect the learning component of the three-chamber assay, in which mice discriminate between a familiar and a novel conspecific. In control animals, this phase typically results in a preference for the novel mouse. In contrast, mice lacking climbing fiber signaling failed to show this preference, consistent with impaired social discrimination or learning. Importantly, our results further suggest that atypical climbing fiber wiring, as seen in our *Pdx1^Cre/+^;vGluT2^fl/fl^*mice, is sufficient to preserve performance in this task, indicating that some intact signaling rather than precise developmental wiring is more important for this form of social behavior.

### Competitive climbing fiber development promotes cerebellar circuit resiliency

We observed a markedly atypical olivocerebellar circuit when climbing fiber neurotransmission was eliminated in only a subset of inputs. Neurotransmission-deficient climbing fibers reduced their Purkinje cell innervation territory by approximately 50%, while competing intact fibers expanded their territory by a similar magnitude. The resulting circuit contains two principal abnormalities: roughly 20–25% of Purkinje cells are innervated by climbing fibers lacking functional neurotransmission, an a comparable fraction is innervated by climbing fibers that would not normally establish synaptic contact in typical development. In total, nearly half of Purkinje cells exhibit atypical climbing fiber connectivity.

Despite this substantial reorganization, behavioral consequences were surprisingly modest. We observed only mild impairments in climbing-fiber-dependent motor assays and no detectable changes in social behaviors. These results suggest that competitive input selection supports the development of cerebellar circuits that are resilient to partial developmental perturbations.

One interpretation is that non-cerebellar circuits involved in motor and social behavior compensate for atypical cerebellar organization. However, such compensation does not imply that the cerebellum is not functionally engaged in these behaviors. Indeed, in the *Ptf1a^Cre/+^;vGluT2^fl/fl^*model, which eliminates climbing fiber neurotransmission more broadly, animals exhibit severe motor deficits and abnormal social behaviors that cannot be rescued by other brain regions. Together, these findings indicate that the relatively mild phenotype observed in partial disruption models likely reflects the robustness of developing circuits to limited perturbations in climbing fiber signaling, rather than redundancy of cerebellar function.

Notably, not all forms of developmental disruption to climbing fibers can be compensated. Prior work in which axon guidance molecules were deleted during early development produced profound miswiring of climbing fibers, preventing decussation and restricting projections to the ipsilateral cerebellar cortex.^17^ This manipulation resulted in severe ataxia and overt motor impairment. Thus, while partial perturbations in neurotransmission can be buffered by developmental mechanisms, more fundamental disruptions in wiring architecture are not tolerated. This suggests that neurotransmission-dependent competition may uniquely support developmental compensation when only a subset of inputs is affected.

Our intersectional genetic mapping further revealed that the *Pdx1;vGluT2* expression domain extends beyond climbing fibers to include additional brainstem and hypothalamic populations. This raises the possibility that some behavioral effects in *Pdx1^Cre/+^;vGluT2^fl/fl^* mice may reflect contributions from these extra-cerebellar circuits. However, the contrast between the modest phenotype in *Pdx1^Cre/+^;vGluT2^fl/fl^*mice and the severe motor and social impairments in *Ptf1a^Cre^;vGluT2^fl/fl^*mice underscores that circuit reorganization within the olivocerebellar system is the dominant determinant of behavioral outcome in the partial disruption model.

The presence of mild motor deficits but intact social behaviors in *Pdx1^Cre/+^;vGluT2^fl/fl^* mice may reflect several, non-mutually exclusive mechanisms. First, altered activity in non-climbing-fiber inferior olive projections to brainstem and spinal circuits may contribute to motor phenotypes independently of cerebellar cortical changes. Second, motor behaviors may require finer circuit precision than the social assays used here. Third, the motor tasks employed may be more sensitive to subtle circuit perturbations than social behavioral paradigms. Finally, motor control may depend more critically on climbing fiber signaling than social behavior, limiting the extent to which developmental compensation can preserve function. Disentangling these possibilities will require future studies using temporally precise, cell-type-specific circuit manipulations.

## Conclusion

In summary, our results provide direct experimental evidence that neurotransmission is essential for climbing fiber competitiveness during Purkinje cell input selection. We further show that climbing fiber neurotransmission is required for a broad range of motor and social behaviors; however, these behaviors are only minimally affected when developmental perturbations are restricted to a subset of climbing fibers.

Together, our findings link competitive climbing fiber development to cerebellum-dependent behavior. We demonstrate that the principles governing input selection enable the formation of resilient cerebellar circuits, even when a substantial fraction of climbing fibers is compromised. Thus, competitive circuit assembly acts as a developmental mechanism that promotes robustness in neural function. Finally, our work shows that atypical circuit organization is not inherently pathological and underscores the importance of caution when inferring functional deficits from structural abnormalities alone.

## METHODS

### Animals

All mice used in the experiments described in this manuscript were housed in an AALAS-certified facility. All studies involving mice were reviewed and approved by the Institutional Animal Care and Use Committee (IACUC) at Virginia Tech (VT). We used the following transgenic mouse lines: *Slc17a6^fl^* (*Vglut2^fl^*; JAX:012898),^52^ Ai14 (*Rosa^lsl-tdTomato^*; JAX: 007914),^53^ Ai65 (*Rosa^ds-tdTomato^*; JAX: 021875),^54^ *Slc17a6^IRES-FlpO^ (Vglut2^FlpO^*, JAX: 030212),^55^ *Ptf1a^Cre^*,^56^ and *Pdx1^Cre^*.^27^ Ear and tail clippings from pre-weaned pups were collected for genotyping and identification of transgenic alleles. Experiments included mice of both sexes, with the day of birth designated as postnatal day 0 (P0). The mice were maintained under a 12-hour light/dark cycle, with environmental conditions set to a temperature range of 68–79°F and humidity levels of 30–50%.

### Tissue processing

Brain tissue was collected for analyses as previously described.^42,57^ Mice were heavily anesthetized with isoflurane and tested for effective sedation by pinching the toe or tail. The chest cavity was accessed, and the heart was penetrated with a butterfly needle. Mice were transcardially perfused with cold phosphate-buffered saline (PBS, 1X) to flush out the blood, then with 4% paraformaldehyde (PFA) to fix the tissue. Tissue was considered fixed by stiffness in the tail and hind paws. Once fixed, mice were decapitated, and the brain was dissected from the skull. Once removed, brains were post-fixed in 4% PFA overnight at 4°C. They were then cryoprotected in a sucrose gradient at 4°C, starting at 10% sucrose in PBS, followed by 20% sucrose in PBS, and finishing in 30% sucrose in PBS. Finally, the tissue was embedded in an optimal cutting temperature (OCT) solution and stored at -80°C. Brain sections were cut sagittally (for climbing fiber counts) or coronally (for inferior olive counts) on a cryostat into 40 µm free-floating tissue sections, which were stored in a PBS-sodium azide bath at 4°C until used for immunohistochemistry. Tissue with tdTomato expression was stored in opaque, light-tight containers at all steps to deter photobleaching.

### Immunohistochemistry

Free-floating brain sections received the following immunofluorescent staining protocol, as previously outlined.^42,57^ Tissue sections were blocked for one hour in 10% normal goat serum with 0.1% Triton-X in phosphate-buffered saline (PBS-T), and then all slices were incubated overnight at room temperature in guinea-pig-anti-Calbindin (1:500; Synaptic Systems; #214004), rabbit-anti-vGluT2 (1:500; Synaptic Systems; 135402), and rabbit-anti-CART (1:2000; Phoenix Pharmaceuticals; H-003-62) diluted in 500uL blocking solution. After the tissue sections were washed three times in PBS-T for five minutes each, the third wash was replaced with 500uL PBS-T with AlexaFluor™ 647 goat-anti-guinea-pig (1:2000; Invitrogen, #A21450), AlexaFluor™ 488 goat-anti-rabbit (1:2000; Invitrogen, #A32731), and DAPI (1:1000), incubated for two hours at room temperature. The tissue sections were washed another three times in PBS-T for five minutes. Finally, the tissue was mounted on glass slides with VECTASHIELD® Plus mounting medium (Vector Laboratories; #H1900). Once dry, the edges of the coverslips were lined with clear acrylic nail polish. Plates containing tissue with tdTomato expression were enclosed completely in aluminum foil wrap during incubation periods to limit photobleaching, and all mounted slides were stored in opaque containers.

### Microscopy and image processing

Photomicrographs of whole mount tissue sections were captured at 20X resolution on a Leica Mica microscope, equipped with automatic stitching software. Images were processed with Leica thundering image processing. Images were processed with the Leica THUNDER software to enhance image clarity. Images’ color brightness and balance were attuned using ImageJ software and cropped to the appropriate size in Adobe Illustrator.

### Image quantification

#### Climbing fiber innervation proportion

We stained sagittally cut tissue sections obtained from P6, P9, P12, P18, P24, or >P56 control or conditional mice with anti-Calbindin and anti-VGlut2 antibodies as previously described.^42,57^ Next, we imaged several tissue sections from 3-5 mice per age and genotype. We imported tissue images into ImageJ and analyzed their molecular layers, where climbing fibers innervate Purkinje cells, as identified by the Calbindin signal. We adjusted the brightness and contrast in the Calbindin, tdTomato, vGluT2, and/or CART channel such that individual Purkinje cells and climbing fibers were visible. We then counted the number of Purkinje cells innervated and not innervated by tdTomato(+), vGluT2(+), or CART(+) climbing fibers. Finally, we compared among slices the mean proportions of Purkinje cells innervated by tdTomato(+), vGluT2(+), or CART(+) fibers and observed differences in the prevalence of tdTomato(+), vGluT2(+), or CART(+) innervation throughout development between control and mutant mice. Final statistics were performed on the average of 3-8 cerebellar sections for each mouse.

#### Relative climbing fiber height

Climbing fiber innervation height was quantified by comparing the extent of individual climbing fiber inputs to the total height of the corresponding Purkinje cell dendrite. Purkinje cell dendrite height was measured from the base of the dendrite (end of the soma) to the boundary of the molecular layer, which represents the full span of the dendrite. The height of the climbing fiber was determined by measuring from the same dendritic base to the most distant detectable tdTomato, vGluT2, or CART signal along the Purkinje cell dendrite. Our final climbing fiber innervation height was then calculated as the ratio of climbing fiber height to total Purkinje cell dendrite height. Final statistics were performed on the average of 3-10 climbing fibers per sagittal section, and 3-6 cerebellar sections for each mouse.

#### Inferior olive neuron proportion

We stained coronally cut tissue sections obtained from P6, P9, P12, P18, or P24 control and conditional mice with anti-Calbindin antibodies as previously described. Next, we imaged several tissue sections from 3-5 mice from each genotype. We imported tissue images into ImageJ and identified the inferior olive as identified by the Calbindin signal. We adjusted the brightness and contrast in the Calbindin and tdTomato channels such that individual inferior olive neurons were visible. We then counted the number of Calbindin(+) inferior olive neurons with and without tdTomato(+). Final statistics were performed on the average from 5-10 inferior olive containing sections for each mouse.

### Electrophysiology

#### Craniotomy for Purkinje cell recordings in head-fixed, anesthetized mice

A craniotomy was performed prior to recordings. Mice were placed in an anesthesia induction chamber with an oxygen flow of 1.0 L/min and a 3.0% isoflurane concentration. The mice were in the chamber for a minimum of 5 minutes, or until they became fully anesthetized. After their breathing went from short, rapid breaths to more regular breathing, the mice were moved from the chamber to a stereotaxic surgery area and placed on a heating pad set to 33°C. Here, the mice were placed into a nose cone to maintain a constant isoflurane flow and were head-fixed using ear bars. Once secured into the surgery area, the isoflurane was lowered to a maintenance level of 0.7 L/Min oxygen flow and 2.0% isoflurane concentration. A toe pinch was done to ensure the mice were still fully anesthetized. If there was a reaction to the toe pinch, then the isoflurane concentration and oxygen flow were increased. Depilatory cream (Nair) was used to remove the fur on the mouse’s head, and an incision was made from the base of the neck to past Bregma. Neck fat was removed from over the cerebellar cortex. Using a microdrill, a craniotomy was performed over the interposed nucleus, 6.3mm posteriorly and 1.25-1.75mm laterally from Bregma.

#### In vivo electrophysiology Purkinje Cell recordings in anesthetized, head-fixed mice

We collected *in vivo*, whole-cell extracellular Purkinje cell recordings in head-fixed anesthetized mice. The isoflurane was set to a lower maintenance level of 0.3 L/Min oxygen flow and 1.0% isoflurane concentration for the recordings. We used TREC microelectrodes (5-8 MΩ, Thomas Recordings) that were attached to an amplifier head-stage (ALA Scientific). A micromanipulator (MP-225A, Sutter Instrument) was used to move the electrode to the recording coordinates that were 6.3mm posteriorly and 1.25-1.75mm laterally from Bregma. Multiple penetrations were made within the lateral coordinate range. Each was 125-150mm apart. An extracellular amplifier (EXT-02B, ALA Scientific) amplified and bandpass filtered (0.3-20 kHz) neural signals. A CED board then digitized the signals. Spike2 software (CED) was used to record neural signals. Purkinje cells were identified based on the presence of complex spikes. Because not all Purkinje cells produce complex spikes in the experimental mouse model *(Pdx1^cre/+^; Vglut2^fl/fl^)*, Purkinje cells were identified based on mean firing rate, spike characteristics, and recording depth (0-3mm). When a Purkinje cell was identified, a 100-second recording was collected. Once a recording session had ended, the mouse was promptly euthanized using an overdose of isoflurane and a secondary method of euthanasia.

#### Spike sorting and analyses of in vivo Purkinje cell recordings

Spike2 software was used to spike sort all 100-s Purkinje cell recordings. Spikes were identified as either simple spikes or complex spikes, and background electrical noise was marked to be excluded from analyses. Following spike sorting, the recordings were run through a custom MATLAB code to calculate parameters measuring spike pattern (neuron) activity.^57,58^ GraphPad Prism (v10.5.0, GraphPad Software, San Diego, CA) was used to run nested T-tests with the parameters for each cell as an independent measurement and mouse ID as the nested variable. We defined our firing pattern parameters as followed: firing rate = total spikes / recording duration; CV = mean (ISI) / stdev (ISI); CV2 = mean (2*(ISI_n_ – ISI_n+1_) / (ISI_n_ + ISI_n+1_)), where ISI = inter spike interval.

### Behavioral analyses

#### Surface righting reflex

The surface righting reflex was tested at P7, P9, P11, and P13.^41^ Pups were aligned in a dorsal position in a clean, empty, flat, solid cage lid. We measured the time until the pups turned onto all four paws. The trial was suspended if the pup did not turn in 60 seconds. The pups were tested three times at each time point.

#### Negative geotaxis reflex

The negative geotaxis reflex was tested at P7, P9, P11, and P13.^41^ Pups were placed facing downward on an inclined surface (∼35°) covered with an absorbent underpad barrier (VWR catalog # 82020-845). We measured the time until the pup turned 90°. The trial was suspended if the pup did not turn in 60 seconds or if it fell down the slope. The pups were tested three times at each time point.

#### Ultrasonic vocalizations – pup

Ultrasonic vocalizations were measured in a social isolation task.^41^ Pups were tested at P7, P9, P11, and P13. Each pup was tested once for 120 s at each time point immediately after being removed from the nest and dam. The mice were placed and monitored in a sound-isolated, ventilated, and light-controlled chamber (Metris SmartChamber), with light and ventilation activated during each recording. Vocalizations were recorded using the Gold Foil Electrostatic Transducer microphone in aluminum shielded housing with a screw connector for tripods and two through-holes for mounting purposes (Metris). Sound was collected, digitized, and analyzed using Metris Sonotrack at a 250 kHz sampling rate via one channel. We recorded the total number of calls.

#### Open Field Assay

The open field assay evaluated motor behavioral characteristics in adult mice tested at 6 weeks of age.^42^ Before each trial, the rectangular open-field apparatus was thoroughly cleaned with 70% ethanol to remove residual odors that could influence motor behavior. Each mouse was individually placed in the center of the apparatus and allowed to explore freely for 10 minutes. The assay was conducted under consistent light conditions and kept free of noise, to ensure that all mice were exposed to the same conditions. Behavioral movements were recorded and analyzed using ANY-maze software, which measured total distance traveled (m) and average speed (m/s) as primary locomotor metrics.

#### Rotarod

The rotarod assessed motor control and coordination of adult mice tested at 6 weeks of age.^42^ Mice were placed on an accelerating rod (4 – 40 rpm over 5 minutes) and required to maintain locomotion in response to increasing speed. The trial ended when a mouse fell off the rod, completed two consecutive rotations (hanging on without walking), or successfully remained on the rod for the full 5 minutes. If the mouse was unable to balance on the rotarod before the trial began, they were marked with an “F” to indicate a failed trial. Each mouse underwent three trials per day for three consecutive days, with a 15-minute rest period in between trials.

#### Three Chamber Assay

The three-chamber assay was employed to evaluate the sociability of adult mice tested after 10 weeks of age.^42,59^ The apparatus consisted of a rectangular, coverless structure divided into three equally sized, opaque compartments. The room was brightly lit with no noise or visual distractions. To acclimate the mice to the environment and the experimenter, they were brought to the behavioral room at least once before the test day by the same individual conducting the experiment. Each mouse was placed in the apparatus for 6 minutes where they were free to explore all sides. On test day, the apparatus and cups (used to place the novelty mice) were sanitized with 70% ethanol to eliminate residual odors prior to the test. We used ANY-maze tracking software to create a protocol for tracking the mouse’s movement, measuring the amount of time each mouse interacted with each zone, and controlling the time in the three stages of the experiment. During the first stage of the experiment, each mouse was first placed in the middle of the apparatus. Once ANY-maze tracked the mouse, we manually lifted the doors open for the test mouse to interact with all sides. There were no social stimuli on either side during this phase. For each mouse to move on to the next stage, they would need to show no side preference by having less than a 50-second difference on each side. If they showed a side preference, they were given four chances to habituate but were excluded from the experiment if they failed the habituation phase. In the second stage, a novel mouse was placed in the cage on one side while the cage on the other side remained empty. Each mouse was again placed in the middle for ANY-maze tracking before lifting the doors for the mouse to explore. In the third stage, the novel mouse from the previous stage becomes the familiar mouse and is switched to the opposite chamber. A new, novel mouse is placed in the other chamber. Each experimental mouse was again placed in the middle for ANY-maze tracking before lifting the doors for the mouse to explore. Each phase was tested for 10 minutes where we measured the time each test mouse interacted in the small zone of the cages. After the last phase was completed, the apparatus and cages were sanitized with 70% ethanol to prepare for the next mouse.

#### Ultrasonic vocalizations – adult

Ultrasonic vocalizations were measured in a courtship experiment.^44,45^ Adult male mice were tested after 10 weeks of age. Before testing, each male test mouse was placed in an empty cage (sterilized with 70% ethanol) with no bedding. The cage was placed in a sound-isolated, ventilated, and light-controlled chamber (Metris SmartChamber) for 8 minutes to habituate the mouse to the testing environment. During testing, each male mouse was tested in two trials, each lasting 120 seconds, where it was paired with a different female mouse of an age at most 3 weeks apart from that of the male. The cage containing the male and female mice was placed in the Metris SmartChamber, with light and ventilation activated during each recording. Vocalizations were recorded using the Gold Foil Electrostatic Transducer microphone in aluminum shielded housing with a screw connector for tripods and two through-holes for mounting purposes (Metris). Sound was collected, digitized, and analyzed using Metris Sonotrack at a 250 kHz sampling rate via one channel. We recorded the total number of calls and reported the average between the two trials.

#### String Pull Assay

The string pull assay assessed the fine motor coordination of adult mice tested after 6 weeks of age. Mice were placed on a reduced food diet of 2g ground chow for females or 2.5g for males for four days before testing. As previously described, on the day before testing, mice were placed in an empty cage with 20 strings of lengths varying 30-100cm draped over the walls, their ends just touching the floor.^60,61^ Attached to half of these strings was a fruit loop on the far end, and the mice were removed after they had pulled 10 strings into the cage or one hour had passed. On testing day, mice were placed in a clear cylindrical apparatus containing one 100cm string with a fruit loop attached to the far end draped over the wall as before. Three 20 min trials were recorded on video per mouse that pulled the string and collected the fruit loop, and only mice that performed in the test were included. Periods of inactivity were trimmed from each trial video, which was slowed to 0.1X speed to quantify normal pulls and three types of pulling errors.^60^ Error rates were calculated from the number of errors per total number of pulls. All mice performed 2 or 3 trials of the assay.

### Statistical analysis

All statistical analysis was performed using GraphPad Prism (v10.5.0, GraphPad Software, San Diego, CA). All statistical tests, comparison groups, significance analyses, and other relevant information for data comparison are specified in Supplementary Table 1. Multiple time-point comparisons were conducted using two-way ANOVA (time-point*group) or repeated measure ANOVA (time-point*group), for the neuroanatomical and behavioral experiments, respectively. We used Tukey post-hoc for post-hoc analyses. Comparisons between two groups at a single timepoint were analyzed using unpaired two-tailed t-tests. Statistical significance was defined as p < 0.05. Effect sizes reported in Table 1 were calculated using GraphPad Prism. R^2^ values were reported for ANOVA analyses, while Cohen’s d was used to determine effect sizes for unpaired t-tests.

## ACKNOWLEDGEMENTS

This work was supported by start-up funds provided by Virginia Tech and the Red Gates Foundation, and NIH-NINDS R00NS13046 to MEvdH. We thank members of the Van der Heijden lab and Dr. Anthony LaMantia for their feedback on this project and the manuscript.

## AUTHOR CONTRIBUTIONS

Conceptualization: JAC, KMC, AML, AEW, MEvdH; Data curation: JAC, KMC, AML, AEW, PAP, JRF, BLD, EAB, ALF, MEvdH; Formal analysis: JAC, AML, MEvdH; Funding acquisition: MEvdH; Investigation: JAC, KMC, AML, AEW, PAP, JRF, BLD, EAB, ALF; Methodology: JAC, KMC, AML, AEW, PAP, JRF, BLD, EAB, ALF, MEvdH; Project administration: JAC, KMC, AML, AEW, MEvdH; Resources: MEvdH; Software: NA; Supervision: MEvdH; Validation: JAC, KMC, AML, AEW, PAP, JRF, BLD, EAB, ALF, MEvdH; Visualization: JAC, KMC, AML, AEW, PAP MEvdH; Writing – original draft: JAC, KMC, AML, AEW, MEvdH; Writing – review & editing: JAC, KMC, AML, AEW, PAP, JRF, BLD, EAB, ALF, MEvdH.

## Notes

### Competing Interest Statement

The authors have declared no competing interest.

