## Supplemental Figure and Table for "Competitive Olivocerebellar Input Selection Promotes Resilient Circuit Formation"

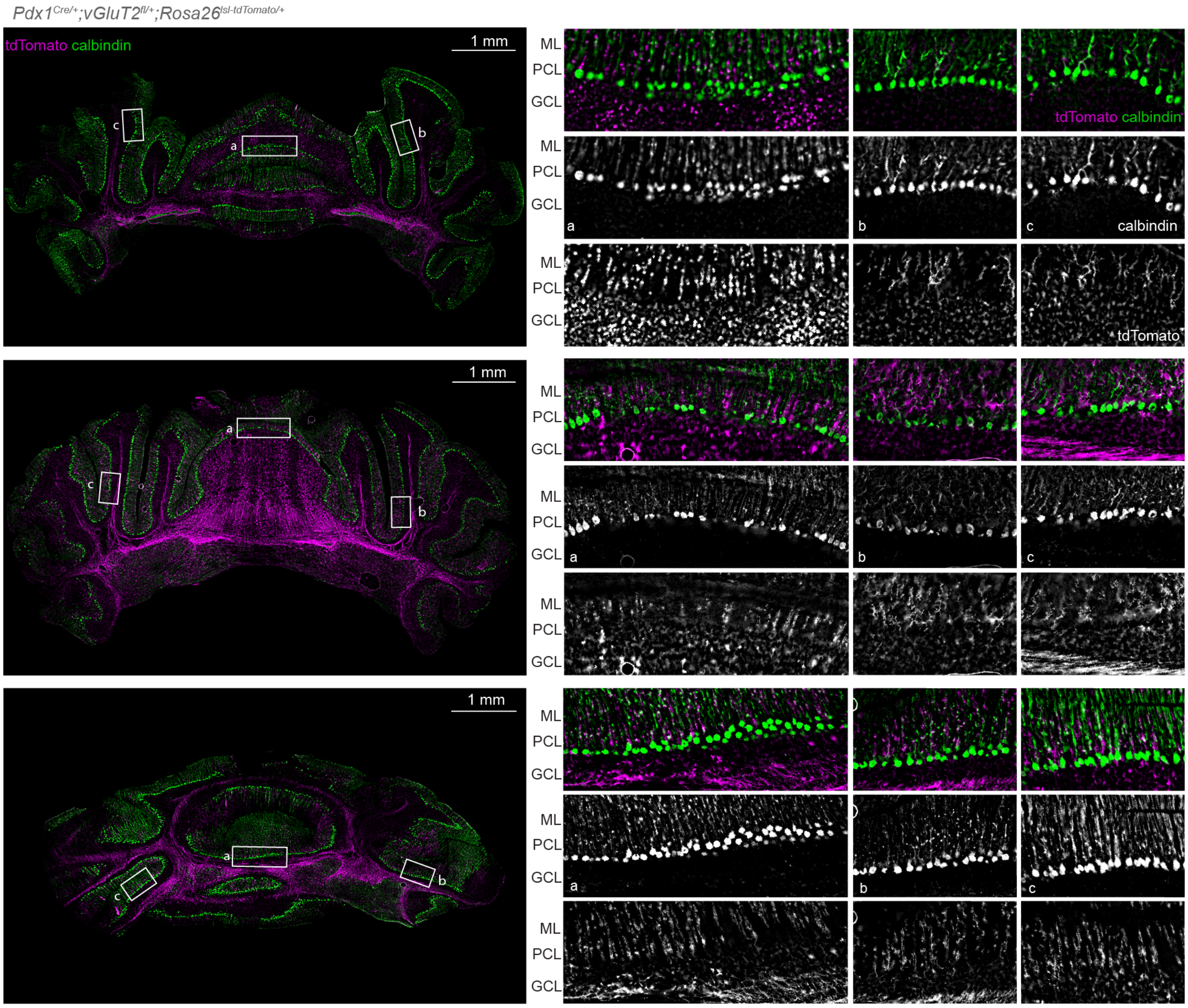


**Supplemental figure 1. tdTomato(+) climbing fibers are present throughout the mediolateral and rostro-caudal cerebellum without apparent patterning.** Robust tdTomato labeling is observed throughout the mediolateral and rostrocaudal coronal sections of the cerebellum, indicating that *Pdx1*$^{Cre/+}$ is expressed in olivo-cerebellar fibers throughout the cerebellar circuitry. Green = calbindin (also in white, second row); Purple = tdTomato (also in white, third row).

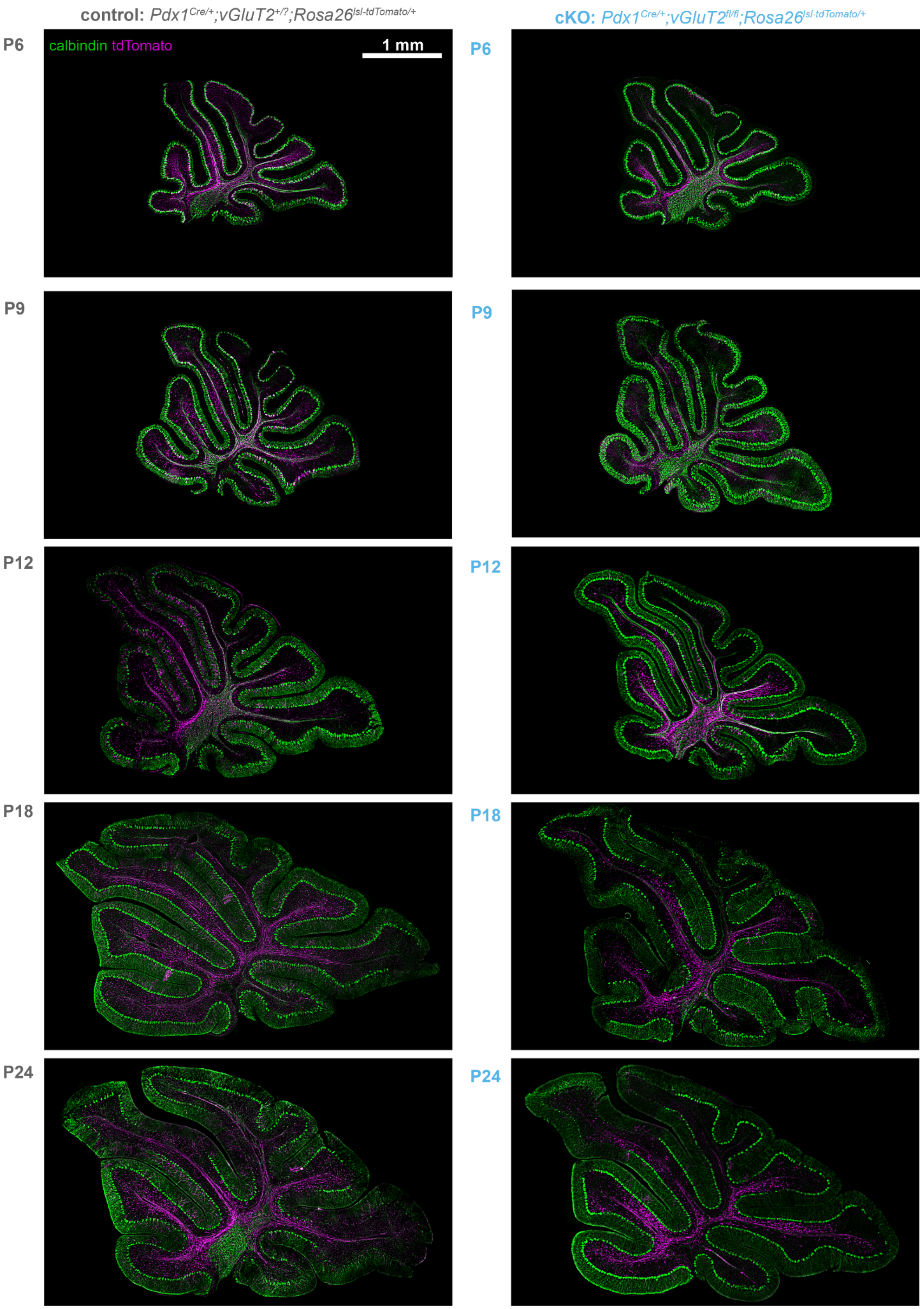


**Supplemental figure 2. Conditional deletion of vGluT2 does not change global cerebellar morphology.** Representative images of the sagittal cerebellar sections from control (*Pdx1$^{Cre}$;vGluT2$^{+/?}$*) and cKO

(*Pdx1*$^{Cre}$*;vGluT2*$^{fl/fl}$) showing comparable anatomical organization. Across all examined timepoints, no obvious differences in regional morphology were observed resulting from vGluT2 deletion. Purple = tdTomato; Green = calbindin.

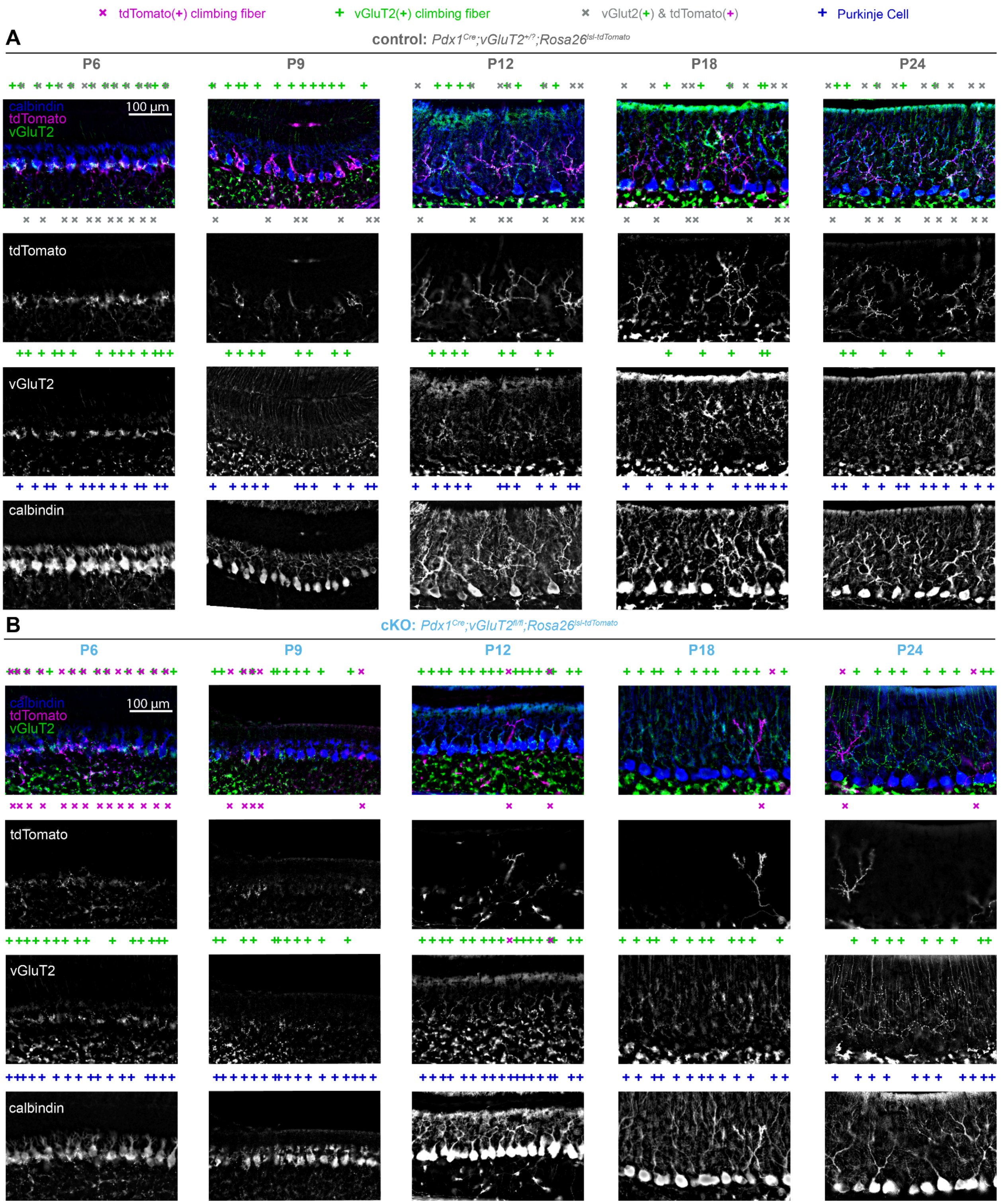


**Supplemental figure 3. tdTomato labeling reveals reduced competitive selection of vGluT2(-) Pdx1Cre/+ climbing fibers during dendritic translocation. A.** At P6, climbing fibers exhibit multi-innervation in the Purkinje cell somata. By P9, climbing fibers undergo dendritic translocation where the “winner” climbing fiber begins to move from the soma to the dendrites. At P12, competitive selection occurs between vGluT2(+) climbing fibers and vGluT2(+) & tdTomato(+) climbing fibers. Following P12, a single climbing fiber is selected to mono-innervate the Purkinje cell dendrite, accompanied by increased dendritic extension and expansion of the molecular layer. Blue = calbindin (Purkinje cells); Purple = tdTomato; Green = vGluT2. **B.** At P6, climbing fibers exhibit multi-innervation in the Purkinje cell somata. By P9, the “winner” climbing fiber begins to translocate from the soma to the dendrites in an activity-dependent manner. At P12,the majority of tdTomato(+) climbing fibers are selectively pruned, while vGluT2(+) climbing fibers are competitively selected to mono-innervate the Purkinje cell dendrite. Blue = calbindin (Purkinje cells); Purple = tdTomato; Green = vGluT2.

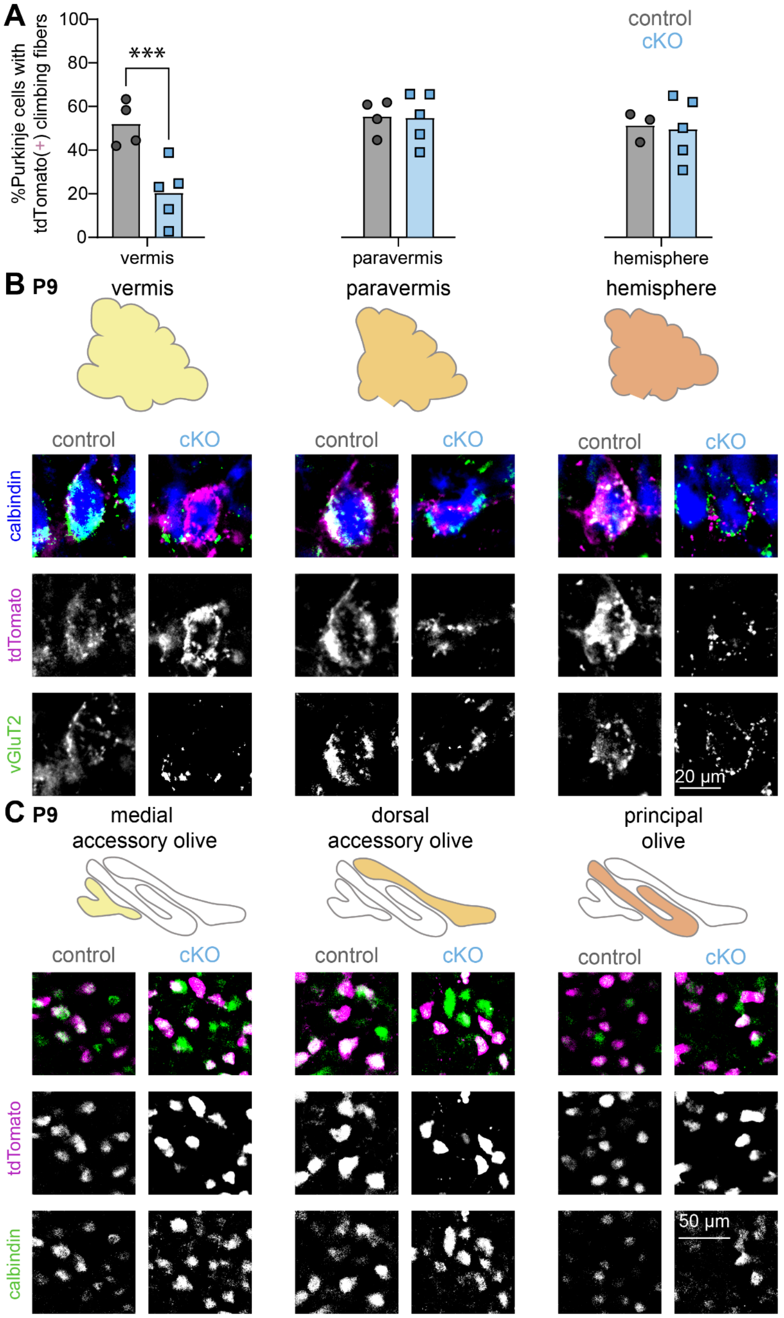


**Supplemental figure 4. Mono and multi-innervation in the same brain at P9 without inferior olive loss in conditional knockout mice. A.** Though paravermal and hemispheric sections show no significant difference in tdTomato(+) innervation between groups, fewer tdTomato(+) climbing fibers are observed in vermal sections from *Pdx1$^{Cre}$;vGluT2$^{fl/fl}$* mice than controls at P9. Vermis: p = 0.0006; paravermis: p = 0.9372; hemisphere p = 0.8435. **B.** Representative Purkinje cells from vermal sections exemplify completed climbing fiber selection in both controls and *Pdx1$^{Cre}$;vGluT2$^{fl/fl}$* mice. While controls similarly show completed climbing fiber selection in paravermal and hemispheric sections, intermingled tdTomato(+) and tdTomato(-) fibers on Purkinje somas reveal multi-innervation remains in these regions in P9 *Pdx1$^{Cre}$;vGluT2$^{fl/fl}$* mice. Blue = calbindin (Purkinje cells); Purple = tdTomato; Green = vGluT2. **C.** Representative images from inferior olive regions that innervate each mediolateral region of the cerebellum show a robust presence of tdTomato(+) neurons innervating the vermis, paravermis, and hemisphere alike in both controls and *Pdx1$^{Cre}$;vGluT2$^{fl/fl}$* mice. Purple = tdTomato; Green = calbindin. All images were obtained from the same animal. Purkinje cell images are representative for 4 out of 5 *Pdx1$^{Cre}$;vGluT2$^{fl/fl}$* mice, and inferior olive neuron images are representative for 3 out of 4 *Pdx1$^{Cre}$;vGluT2$^{fl/fl}$* mice.

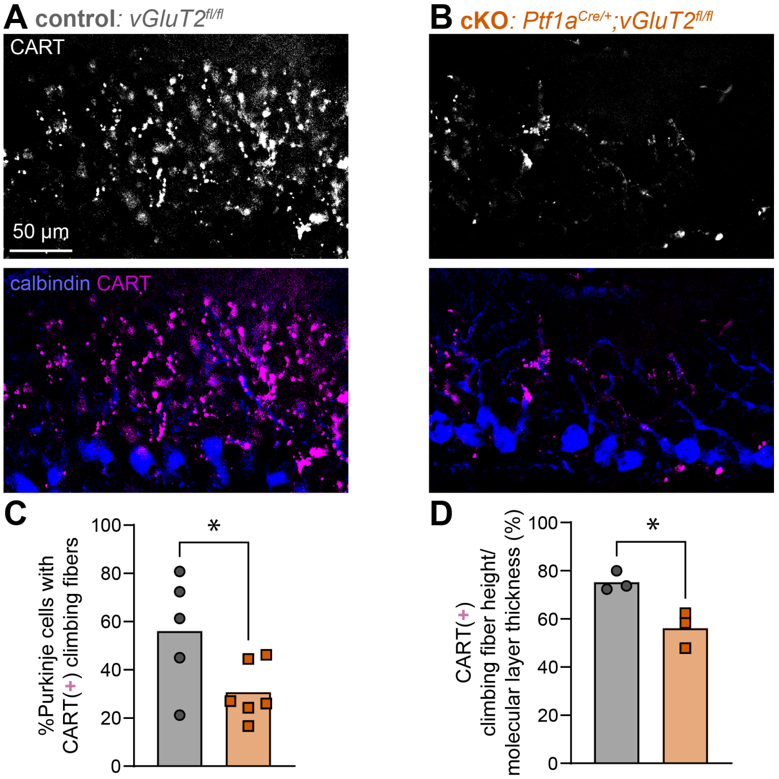


**Supplemental figure 5. CART labeling reveals reduced competitive selection of vGluT2(-) climbing fibers despite sparse vGluT2(+) input. A.** Representative image shows abundant CART (purple) labeling of climbing fibers and calbindin (blue) labeling of Purkinje cells in control mice. **B.** Representative image shows sparse CART (purple) labeling of climbing fibers and calbindin (blue) labeling of Purkinje cells in *Ptf1a$^{Cre/+}$;vGluT2$^{fl/fl}$* mice. **C.** Quantification confirmed a reduction of CART signal in the *Ptf1a$^{Cre/+}$;vGluT2$^{fl/fl}$* cerebella (p = 0.0461; control: N=5 mice, n=25 sections , *n*=1,073 Purkinje cells; cKO: N=6 mice, n=30 sections, *n*=1128). **D.** Quantification confirmed a reduction in climbing fiber height in the *Ptf1a$^{Cre/+}$;vGluT2$^{fl/fl}$* cerebella (p = 0.0177; N=3 mice, n=15 sections , *n*=150 Purkinje cells; cKO: N=3 mice, n=15 sections, *n*=150).

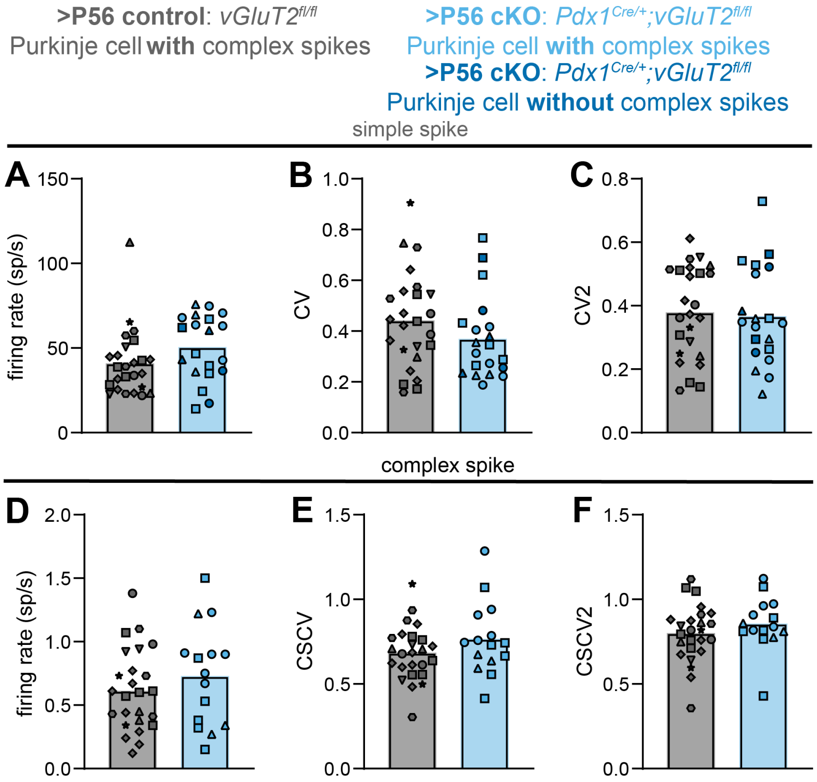


**Supplemental figure 6. No difference in Purkinje cell simple and complex spike firing patterns between control and *Pdx1$^{Cre}$;vGluT2$^{fl/fl}$* mice. A.** No differences were found in the SS firing rate (p = 0.1402, d = 0.25), **B.** SS CV (spike pattern, p = 0.2435, d = 0.17), **C.** SS CV2 (spike regularity, p = 0.8024, d = 0.01). **D.** No differences were found in the CS firing rate (p = 0.6287, d = 0.03), **E.** CS CV (spike pattern, p = 0.2364, d = 0.17), **F.** CS CV2 (spike regularity, p = 0.3227, d = 0.12). Data points from control mice in gray, *Pdx1$^{Cre}$;vGluT2$^{fl/fl}$* mice (with complex spikes) in light blue, and *Pdx1$^{Cre}$;vGluT2$^{fl/fl}$* mice (without complex spikes) in dark blue. Each shape represents a different mouse. Control: N = 7 mice, n = 25 cells; *Pdx1$^{Cre}$;vGluT2$^{fl/fl}$*: N = 3 mice, n = 20 cells (simple spikes), n = 15 cells (complex spikes).

**Supplemental Table 1. Information about statistical analyses.**

Results from statistical analysis. For T-tests we included the t-statistics and for ANOVAs we reported the F-statistics. We reported the eta-square for all t-tests and F-tests except for the nested T-tests where we reported Cohen's d. Statistically significant differences are bolded and highlighted in peach, middle and large effect sizes are highlighted in light and deep yellow, respectively.

| | Figure panel | Test | Measurement | Comparison groups | Genotype / Age / Interaction | F(DFn, DFd) = ... / t(df) = ... | P = ... | eta-square = ... / cohen's d = ... |
|---|---|---|---|---|---|---|---|---|
| tdTomato(+) climbing fibers | 2F | ANOVA | % Purkinje cells with tdTomato(+) climbing fibers | Control vs. Pdx1-cKO | Age x Genotype | F (4, 28) = 6.397 | **P=0.0009** | 0.119 |
| | | | | | Age | F (4, 28) = 31.60 | **P<0.0001** | 0.588 |
| | | | | | Genotype | F (1, 28) = 48.01 | **P<0.0001** | 0.2233 |
| | 2G | T-test | % Purkinje cells with **only** tdTomato (+) climbing fibers | Control vs. Pdx1-cKO | Genotype | t(5) = 1.309 | P=0.2720 | 0.2251 |
| | 2K | T-test | tdTomato(+) climbing fiber height/ molecular layer thickness (%) | Control vs. Pdx1-cKO | Genotype | t(5) = 5.029 | **P=0.0040** | 0.8507 |
| | 2L | T-test | vGluT2(+) climbing fiber height/ molecular layer thickness (%) | Control vs. Pdx1-cKO | Genotype | t(5) = 0.5729 | P=0.5915 | 0.0616 |
| tdTomato(+) inferior olive neurons | 3C | ANOVA | % tdTomato(+) inferior olive neurons | Control vs. Pdx1-cKO | Age x Genotype | F (4, 28) = 2.103 | P=0.1070 | 0.1136903394 |
| | | | | | Age | F (4, 28) = 8.767 | **P=0.0001** | 0.4736842105 |
| | | | | | Genotype | F (1, 28) = 30.55 | **P<0.0001** | 0.4126254501 |
| | 3D | T-test | %tdTomato(+) in cKO normalized to control | climbing fiber vs. inferior olive | Cell type | t(21) = 3.197 | **P=0.0043** | 0.3274 |
| vGlut2(+) climbing fibers | 4G | ANOVA | %Purkinje cells with vGluT2(+) climbing fibers | Control vs. Pdx1-cKO | Age x Genotype | F (1, 10) = 1.424 | P=0.2603 | 0.03747 |
| | | | | | Age | F (1, 10) = 1.012 | P=0.3382 | 0.02662 |
| | | | | | Genotype | F (1, 10) = 27.37 | **P=0.0004** | 0.7201 |
| | 4L | Fisher's Exact Test | %Purkinje cells with complex spike | Control vs. Pdx1-cKO | Genotype | NA | **P=0.0127** | NA |
| motor behaviors | 6C | ANOVA | Surface righting reflex | Control vs. Pdx1-cKO | Age x Genotype | F (3, 189) = 9.119 | **P<0.0001** | 0.06511 |
| | | | | | Age | F (1.169, 73.65) = 24.58 | **P<0.0001** | 0.1755 |
| | | | | | Genotype | F (1, 63) = 10.96 | **P=0.0015** | 0.04723 |
| | 6D | ANOVA | Surface righting reflex | Control vs. Ptf1a-cKO | Age x Genotype | F (3, 150) = 22.15 | **P<0.0001** | 0.1113 |
| | | | | | Age | F (2.345, 117.3) = 35.32 | **P<0.0001** | 0.1775 |
| | | | | | Genotype | F (1, 50) = 70.28 | **P<0.0001** | 0.2621 |
| | 6F | ANOVA | Negative geotaxis reflex | Control vs. Pdx1-cKO | Age x Genotype | F (3, 189) = 0.3882 | P=0.7617 | 0.002983 |
| | | | | | Age | F (2.504, 157.7) = 30.17 | **P<0.0001** | 0.2318 |
| | | | | | Genotype | F (1, 63) = 0.05986 | P=0.8075 | 0.0002651 |
| | 6G | ANOVA | Negative geotaxis reflex | Control vs. Ptf1a-cKO | Age x Genotype | F (3, 150) = 0.7668 | P=0.5143 | 0.002727 |
| | | | | | Age | F (2.423, 121.2) = 10.02 | **P<0.0001** | 0.03562 |
| | | | | | Genotype | F (1, 50) = 413.6 | **P<0.0001** | 0.6993 |
| | 6I | ANOVA | Rotarod | Control vs. Pdx1-cKO | Day x Genotype | F (2, 64) = 1.300 | P=0.2797 | 0.01575 |
| | | | | | Day | F (1.422, 45.51) = 7.169 | **P=0.0049** | 0.08685 |
| | | | | | Genotype | F (1, 32) = 12.03 | **P=0.0015** | 0.1412 |
| | 6J | ANOVA | Rotarod | Control vs. Ptf1a-cKO | Day x Genotype | F (2, 56) = 0.1871 | P=0.8299 | 0.0002077 |
| | | | | | Day | F (1.664, 46.60) = 12.13 | **P=0.0001** | 0.01346 |

| | | | | | | | | |
|---|---|---|---|---|---|---|---|---|
| | | | | | Genotype | F (1, 28) = 309.9 | **P<0.0001** | 0.8758 |
| | 6L | T-test | Open Field - Speed | Control vs. Pdx1-cKO | Genotype | t(32) = 0.2903 | P=0.7735 | 0.002626 |
| | 6M | T-test | Open Field - Distance | Control vs. Pdx1-cKO | Genotype | t(32) = 0.3110 | P=0.7578 | 0.003014 |
| | 6N | T-test | Open Field - Speed | Control vs. Ptf1a-cKO | Genotype | t(28) = 3.965 | **P=0.0005** | 0.3596 |
| | 6O | T-test | Open Field - Distance | Control vs. Ptf1a-cKO | Genotype | t(28) = 3.974 | **P=0.0005** | 0.3607 |
| | 6Q | T-test | Partial miss rate | Control vs. Pdx1-cKO | Genotype | t(26) = 0.8726 | P=0.3909 | 0.02845 |
| | 6R | T-test | Full miss rate | Control vs. Pdx1-cKO | Genotype | t(26) = 2.063 | **P=0.0493** | 0.1406 |
| | 6S | T-test | Double pull rate | Control vs. Pdx1-cKO | Genotype | t(26) = 3.177 | **P=0.0038** | 0.2796 |
| **social behaviors** | 7C | ANOVA | Social Isolation Vocalizations | Control vs. Pdx1-cKO | Region x Genotype | F (3, 189) = 0.7986 | **P=0.0183** | 0.3482949957 |
| | | | | | Region | F (2.451, 154.4) = 8.972 | **P=0.0078** | 0.4428582593 |
| | | | | | Genotype | F (1, 63) = 0.7110 | **P=0.0247** | 0.208846745 |
| | 7D | ANOVA | Social Isolation Vocalizations | Control vs. Ptf1a-cKO | Age x Genotype | F (3, 150) = 2.149 | P=0.0965 | 0.0261 |
| | | | | | Age | F (2.551, 127.6) = 2.111 | P=0.1119 | 0.02564 |
| | | | | | Genotype | F (1, 50) = 4.776 | **P=0.0336** | 0.02964 |
| | 7F | T-test | Mating Vocalizations | Control vs. Pdx1-cKO | Genotype | t(25) = 1.063 | P=0.2979 | 0.04325 |
| | 7G | T-test | Mating Vocalizations | Control vs. Ptf1a-cKO | Genotype | t(21) = 1.703 | P=0.1034 | 0.1213 |
| | 7I | T-test | 3 Chamber: Novel/Empty | Control vs. Pdx1-cKO | Genotype | t(29) = 0.4616 | P=0.6478 | 0.007294 |
| | 7J | T-test | 3 Chamber: Novel/Empty | Control vs. Ptf1a-cKO | Genotype | t(33) = 0.1601 | P=0.1601 | 0.05889 |
| | 7L | T-test | 3 Chamber: Novel/Familiar | Control vs. Pdx1-cKO | Genotype | t(29) = 0.04834 | P=0.9618 | 0.00008059 |
| | 7M | T-test | 3 Chamber: Novel/Familiar | Control vs. Ptf1a-cKO | Genotype | t(33) = 2.562 | **P=0.0152** | 0.1659 |
| **P9 tdTomato(+) climbing fibers** | S4A | ANOVA | % Purkinje cells with tdTomato(+) climbing fibers | Control vs. Pdx1-cKO | Age x Genotype | F (2, 20) = 4.923 | **P=0.0009** | 0.119 |
| | | | | | Age | F (2, 20) = 6.257 | **P<0.0001** | 0.588 |
| | | | | | Genotype | F (1, 20) = 5.902 | **P<0.0001** | 0.2233 |
| **CART(+) climbing fibers** | S5C | T-test | % Purkinje cells with CART(+) climbing fibers | Control vs. Ptf1a-cKO | Genotype | t(9) = 2.311 | **P=0.0461** | 0.3725 |
| | S5D | T-test | CART(+) climbing fiber height/ molecular layer thickness (%) | Control vs. Ptf1a-cKO | Genotype | t(4) = 3.886 | **P=0.0177** | 0.7906 |
| **Purkinje cell spike properties** | S6A | nested T-test | simple spike - firing rate | Control vs. Pdx1-cKO | Genotype | t(8) = 1.638 | P=0.1402 | 0.2440715532 |
| | S6B | nested T-test | simple spike - CV | Control vs. Pdx1-cKO | Genotype | t(8) = 1.259 | P=0.2435 | 0.1876792281 |
| | S6C | nested T-test | simple spike - CV2 | Control vs. Pdx1-cKO | Genotype | t(8) = 0.2587 | P=0.8024 | 0.03857504856 |
| | S6D | nested T-test | complex spike - firing rate | Control vs. Pdx1-cKO | Genotype | t(8) = 0.5027 | P=0.6287 | 0.07946996504 |
| | S6E | nested T-test | complex spike - CV | Control vs. Pdx1-cKO | Genotype | t(8) = 1.639 | P=0.7655 | 0.1908366325 |
| | S6F | nested T-test | complex spike - CV2 | Control vs. Pdx1-cKO | Genotype | t(8) = 1.054 | P=0.3227 | 0.1571253609 |
